# Optimizing checkpoint strategies based on first principles predicts experimental DNA damage checkpoint override times

**DOI:** 10.1101/2020.08.14.251504

**Authors:** Ahmad Sadeghi, Roxane Dervey, Vojislav Gligorovski, Sahand Jamal Rahi

## Abstract

Why biological quality-control systems fail is often mysterious. Specifically, checkpoints such as the DNA damage checkpoint or the spindle assembly checkpoint are overriden after prolonged arrests allowing cells to continue dividing despite the continued presence of errors.^1–4^ Although critical for biological systems, checkpoint override is poorly understood quantitatively by experiment or theory. Override may represent a trade-off between risk and speed, a fundamental principle explaining biological phenomena.^5,6^ Here, we derive the first, general theory of optimal checkpoint strategies, balancing risk and opportunities for growth. We demonstrate that the mathematical problem of finding the optimal strategy maps onto the question of calculating the optimal absorbing boundary for a random walk, which we show can be solved efficiently recursively. The theory predicts the optimal override strategy without any free parameters based on two inputs, the statistics i) of error correction and ii) of survival. We apply the theory to the prominent example of the DNA damage checkpoint in budding yeast (*Saccharomyces cerevisiae*) experimentally. Using a novel fluorescent construct which allowed cells with DNA breaks to be isolated by flow cytometry, we quantified i) the probability distribution function of repair for a double-strand DNA break (DSB), including for the critically important, rare events deep in the tail of the distribution, as well as ii) the survival probability if the checkpoint was overridden. Based on these two measurements, the optimal checkpoint theory predicted remarkably accurately the DNA damage checkpoint override times as a function of DSB numbers, which we measured precisely at the single-cell level. Our multi-DSB results refine well-known bulk culture measurements^7^ and show that override is a more general phenomenon than previously thought. Further, we show for the first time that override is an advantageous strategy in cells with wild-type DNA repair genes. The universal nature of the balance between risk and self-replication opportunity is in principle relevant to many other systems, including other checkpoints, developmental decisions^8^, or reprogramming of cancer cells^9^, suggesting potential further applications of the theory.

## Introduction

Cell cycle checkpoints arrest the self-replication process and increase the chances of survival when a cell encounters errors. ^4,10,11^ However, checkpoints show more complex behaviors than merely gating the cell cycle depending on the presence or absence of errors. After prolonged arrests, checkpoints such as the DNA damage checkpoint or the spindle assembly checkpoint (SAC) are overridden in the continued presence of errors, which is associated with genomic instability and aneuploidy in yeast and mammalian cells. ^3,12–16^ (We opt for the more general term ‘override’ in place of ‘adaptation’, ‘slippage’, or ‘leakage’^3^.)

Checkpoints determine the success rate and the speed of the cell cycle after a cell encounters an error. A risk-speed trade-off emerges generally in biological surveillance systems: Whenever self-replication arrests because of an error, opportunities for producing offspring necessarily decrease because of the delay. On the other hand, delays may be necessary for correcting errors, e.g., for DNA damage repair.^17^

Despite the importance of checkpoints to biology, there are no quantitative theories of checkpoint trade-offs or strategies. Many researchers have pointed out that checkpoints may balance opposing demands on repair success and speed.^7,12,16,18^ However, these ideas have so far not been developed into a quantitative theory of checkpoint strategies, which could be supported by experimental measurements of the parameters of the theory and by validation of new predictions. Chemical kinetic models represent the dynamics of the molecular components of checkpoints^19–21^ but they do not indicate potential strategies for checkpoints in an obvious manner as they primarily describe the state of the system in time. Similarly, the potential balances that checkpoints strike are not straightforwardly explained by speed-accuracy-energy dissipation relations or decision-making frameworks proposed for other systems: Sensing in chemotaxis and mechanisms for kinetic proofreading address the difficulties of ascertaining information and decision-making in the presence of noise.^5,22–37^ This contrasts with the case of checkpoints where errors such as DNA damage or poor chromosome-spindle attachment are signaled reliably;^10^ uncertainty arises from the seemingly stochastic nature of the outcomes (survival, sickness, or death).

Experiments support the existence of a balance between risk and speed qualitatively: Budding yeast cells with dysfunctional DNA damage (rad9Δ) or spindle assembly checkpoints (mad2Δ), which arrest very briefly^4^ or not noticeably^10^, die at much higher rates in the presence of DNA damage or poor spindle-chromosome binding.^10^ Multiple experimental challenges have to be overcome in order to resolve checkpoint strategies more quantitatively:

- The differences between strategies may manifest in rare events, which, by their nature, are difficult to capture experimentally, e.g., rare, late repair events. (Over evolutionary timescales or in large populations, rare beneficial outcomes can nevertheless suffice for selection and fixation.)
- Checkpoints and repair systems are complicated, involving dozens of genes and pathways, which makes it difficult to isolate and study specific effects without confounding factors.

The DNA damage checkpoint in budding yeast, which has been intensively studied by molecular biology and genetics,^4,10^ is particularly ripe for quantitative, system-level insights. In haploid cells, which cannot utilize highly efficient template-based DNA repair mechanisms such as homologous recombination, DSBs remain unrepaired in a substantial fraction of cells for many hours.^38^ For a single unrepaired DSB near the left telomere of chromosome VII, the MAT locus, or URA3, it was observed that after an ≈8 hr arrest at the G2/M checkpoint, cells proceeded with the cell cycle although the DSB was not repaired.^1,2,7^ Difficult-to-quantify but possibly large amounts of DNA damage due to X-rays^12^ or telomere dysfunction^2,15^ also lead to checkpoint override. A number of mutations have been found to prevent or reduce checkpoint override: cdc5-ad (missense mutation in CDC5)^2^, ckb2Δ^2^, yku70Δ^7^, and others^39,40^.

The existence of mutations that abrogate override underscores the possibility that DNA damage checkpoint override is functionally important to yeast cells. However, past work has not shown that override has a benefit for wild-type cells nor explained the importance of its timing (≈8 hrs): A key test to establish whether override has a functional role at all is whether cells with DNA damage produce more offspring on average if they override their checkpoint. This was found not to be the case; in haploid yeast cells, no significant difference was found between survival rates of override-performing (CDC5) or override-unable (cdc5-ad) cells after X-ray irradiation, which causes DSBs.^12^ (A situation in which checkpoint override was beneficial was created by artificially blocking critical pathways for DNA repair (rad52Δ) or telomere maintenance (tlc1Δ).^12,16,18^) Furthermore, it was found that DNA damage checkpoint override occurred only if one endonuclease-generated DSB was present but not with two DSBs^7^, indicating that override is not a universal response to DNA damage in yeast.

A key missing empirical ingredient for a quantitative analysis of checkpoint strategies is statistics on rare repair events. In the absence of a genomic template with a similar DNA sequence, repair occurs by non-homologous end joining (NHEJ)^41^, whose timing and efficiency after DSB induction has been measured, for example, by performing qPCR^42–45^ or other quantification methods^46^ in bulk culture at regular time points after the break event. These measurements showed broadly distributed repair times such that a typical repair timescale could not be easily deduced (≈50% repair in about 6 hrs^45^). Bulk culture assays may be too insensitive to quantify rare and late repair events.

Here, we present

1. the first, general theory of optimal checkpoint strategies, balancing the risk of death and opportunities for growth,
2. measurements of the two parameters of the theory, the probability distribution function of repair and the survival probability after checkpoint override,
3. predictions of the theory: optimal override times as a function of the numbers of DSBs,
4. quantitative experimental verification of the predictions, and
5. the first measurements showing that override is indeed advantageous with wild-type DNA repair machinery.

## Results

### Theoretical results

#### Arrest implies an exponential fitness penalty

To compute the fitness effects of different checkpoint strategies, we compared an arrested cell to other cells in the population. The two canonical scenarios for population dynamics are exponential growth (Fig. 1 A) and stochastic birth-death processes in a fixed-size population (Fig. 1 B).^47^

**Fig. 1:**
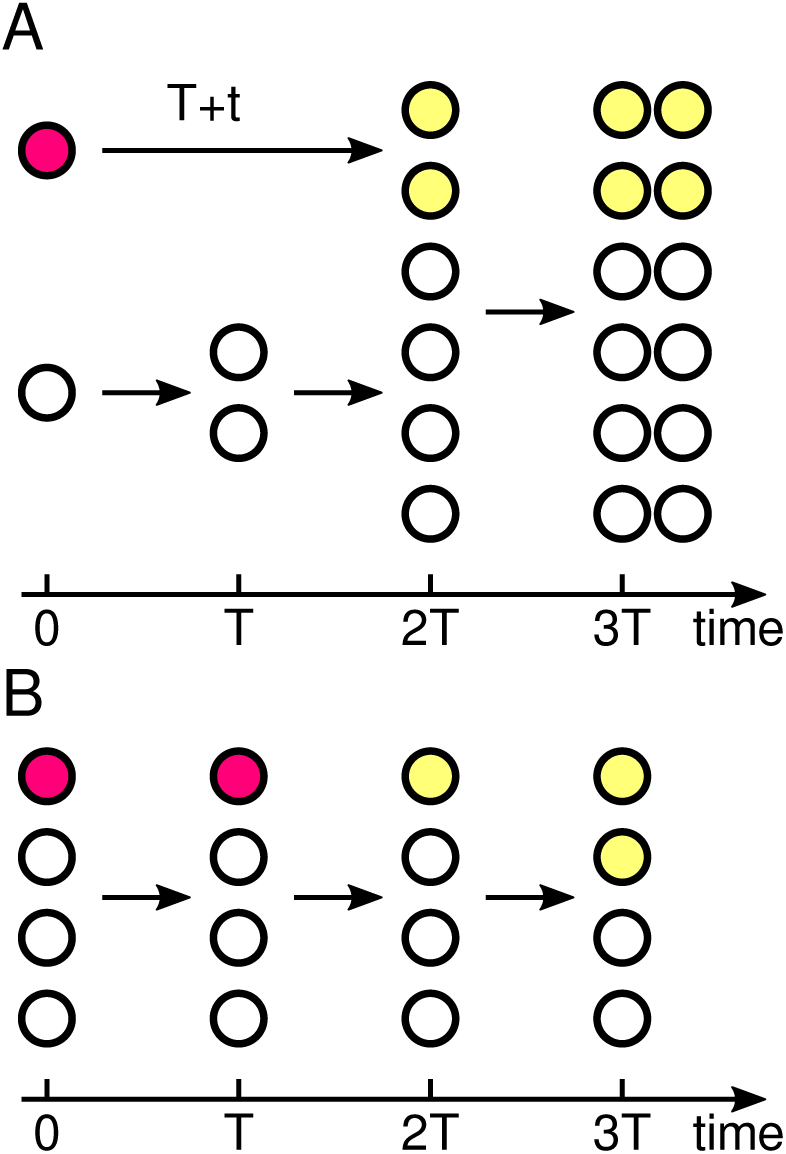
Scenarios for the self-replication dynamics of a checkpoint-arrested cell (magenta circle) in a population of cells (white) with generation time *T*. A: The exponential growth with doubling time *T* is delayed by a checkpoint arrest of duration *t* (= *T* in this illustration) after which both, one, or no progeny (yellow circles) remains viable. Only the case where both progeny are viable is illustrated. B: In a population of fixed size *N*, random birth-death processes change the proportions of different clones, as in the Wright-Fisher or Moran models.

In exponential growth, the expected number of progeny of a checkpoint-arrested cell is reduced by 2^−*t*/*T*^ (*P*_*2*_ + *P*_1_ /2), where *t* is the arrest time, *P*_2_ the probability that both progeny proliferate after the arrest, and *P*_1_ the probability that only one progeny proliferates (Fig. 1 A). (In this presentation, cells either die or produce healthy progeny, in agreement with experimental observations, see Supplementary Information.)

We also analyzed the effect of checkpoint arrest on the expected number of progeny in two classical fixed-size population models, the Wright-Fisher and the Moran models^47^ (Supplementary Information). Here, the reduction in the expected number of progeny has the same form as for exponential growth: the product of a penalty for the arrest time, which is exponential, multiplied by a probability of proliferation. Thus, one can represent the arrest penalty by a general factor *e*^−*t*/*τ*^, which applies in all three cases and where *τ* absorbs model-specific parameters, for example, *τ* = *T* / log 2 for exponential growth.

#### Winning strategy maximizes the arithmetic mean of the future progeny

To evaluate how different strategies influence an organism’s reproductive success, one of two quantities is commonly taken as a fitness function, i) the number of offspring averaged over all possible outcomes or ii) the expected number of offspring in a typical scenario.^5,47,48^ In some instances, such as in fluctuating environments, the typical series of outcomes (ii) is thought to be the more relevant quantity for evolutionary selection^47,48^, favoring bet hedging^49^. However, since errors and checkpoint arrests affect cells presumably randomly, averaging over the predicted number of offspring (i) is the appropriate fitness function to consider here, as we illustrate by contrasting a random checkpoint arrest model with a fluctuating-environment model in the Supplementary Information.

#### Checkpoint strategy parameters

In addition to the arrest penalty, an experimentally accessible set of parameters for analyzing checkpoint strategies are needed:

- The remaining number of ***e***rrors *E* (*t*) in a cell, for example, DSBs (potentially signaled by the amount of RPA covering resected DNA) or unattached kinetochores after arresting for time *t. E*_0_ = *E* (0) denotes the initial number of errors, and *E* (*t*) decreases with the arrest time *t* when errors are corrected.
- The probability *r* ({*t*_*i*_}) that the errors are ***r*** epaired at times 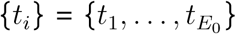. When needed, we will assume that each error is repaired independently with the cumulative probability density function *ρ*(*t*) for the repair of a single error.
- The probability *s* (*t* l {*t*_*i*_}) of ***s***urvival if a checkpoint is overridden after an arrest of duration *t* given that repairs take place at times {*t*_*i*_}. (A repair time *t*_*i*_ after *t* (*t*_*i*_ > *t*) indicates that error *i* was not fixed before the checkpoint is overridden.) To simplify, we will just write *s*(*E*) when the survival probability only depends on the number of remaining errors *E*(*t*). If each error reduces the survival probability by the same amount, we will write *s*(*E*) = *σ*^*E*^.
- The probability of cell cycle ***a*** dvancement *a* (*t* l {*t*_*i*_}), that is, of override if there are unrepaired errors or of continuation of the self-replication cycle if there are no errors. The cell cycle advancement probability *a* (*t* l {*t*_*i*_}) represents the checkpoint strategy and is the function that we seek to compute.

*E* (*t*), *r* (*t* l {*t*_*i*_}), and *s t t*_*i*_ represent the state of the cell and are determined by biochemical processes. The function *a* (*t* l {*t*_*i*_}) represents the behavior of the checkpoint whose effect on the reproductive success of the cell we analyze.

#### General mathematical description of checkpoint strategies

Combining the above probabilities, the fitness coefficient *f* is the factor by which the number of progeny is changed due to the damage and the subsequent behavior of the checkpoint:

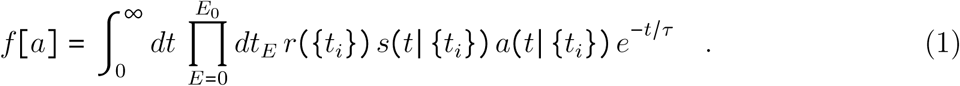

Additional information regarding this expression is given in the Supplementary Information. To visualize the processes that this functional represents, we make two simplifications: each repair occurs independently and each error reduces the survival probability by the same amount. This allows a graphical interpretation of the mathematical expressions, depicted in Fig. 2 A.

**Fig. 2:**
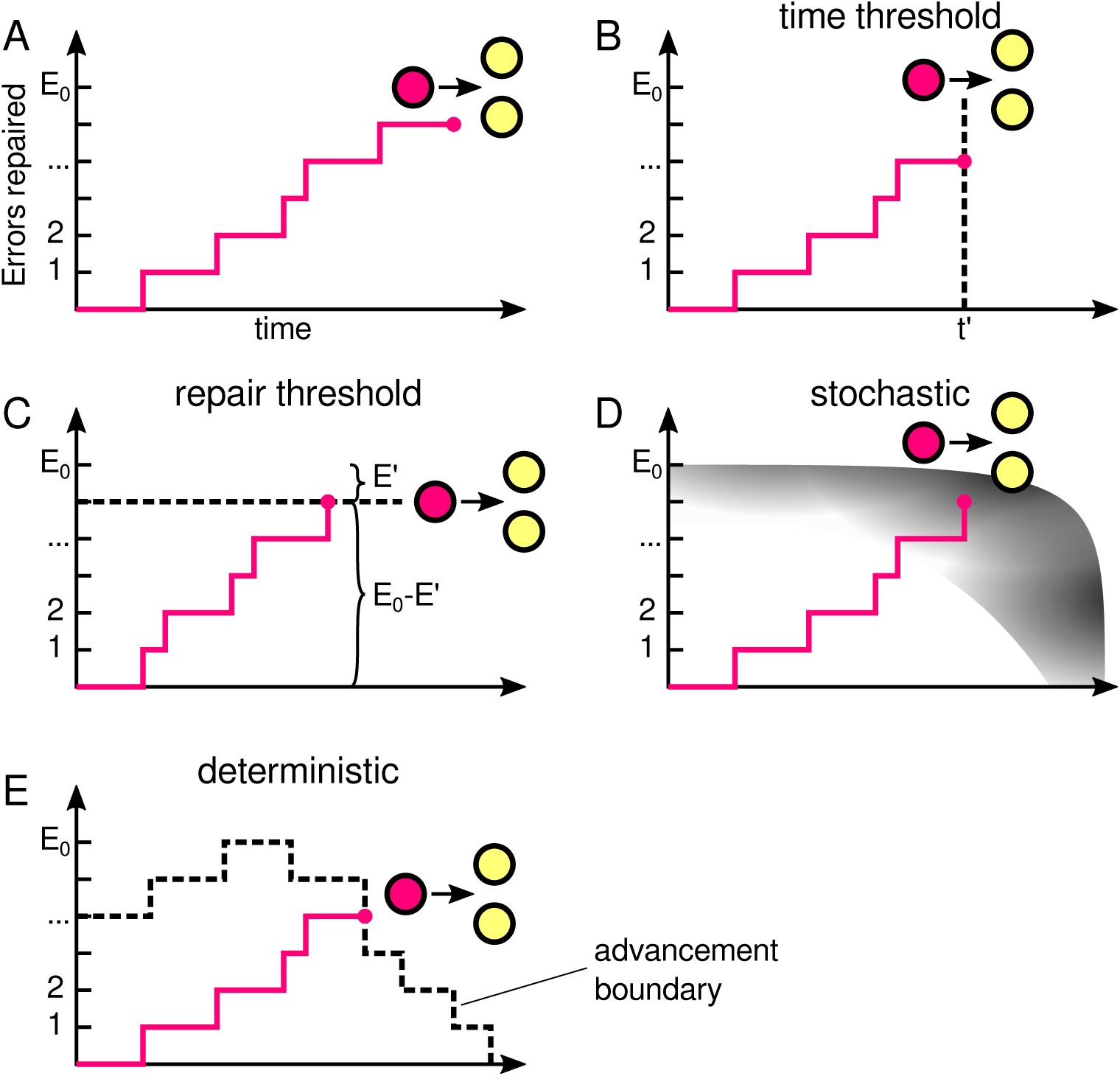
Pictorial representation of checkpoint strategies. A: A checkpoint-arrested cell (red) moves to the right as time increases and moves up as repairs occur probabilistically in accordance with the checkpoint strategy, the cell divides into two cells (yellow). The progeny may or may not be viable. B: In the timer strategy, once a checkpoint is activated, the cell arrests until time *t′* and then divides. The advancement boundary is a vertical line. C: In the repair threshold strategy, cells arrest until *E′* errors remain and then divide. The advancement boundary is a horizontal line. D, E: The optimal strategy could, in principle, be probabilistic (panel D) or deterministic (panel E).

#### Optimal checkpoint strategy

For comparison with the optimal strategy, we also analyzed two other plausible checkpoint strategies, a ‘timer’ override strategy (Fig. 2 B) and an ‘error-threshold’ override strategy (Fig. 2 C), described in the Supplementary Information.

In general, the optimal checkpoint strategy could be stochastic; that is, depending on the state of the cell and on predictions about future outcomes, the optimal decision could be to advance the cell cycle with a probability between zero and one. As explained in the Supplementary Information, we have proven that the optimal strategy cannot be stochastic (illustrated by a gray region in Fig. 2 D) but must be deterministic (dashed lines in Fig. 2 E). This means that the optimal advancement probability *a*(*t*, {*t*_*i*_}) is a Dirac delta (*δ*) function in *t*; the advancement probability can be represented by a sharp decision boundary in the error-time plane (Fig. 2 E), which we refer to as the ‘advancement boundary’. Once a cell reaches this boundary, the optimal decision is to advance the cell cycle.

The problem of solving for the optimal strategy is, in principle, complicated because of the branching nature of the decisions; the consequences of a decision at time *t* and with *E* (*t*) remaining errors depend on an infinite number of possible future events. However, the optimal checkpoint strategy can be solved recursively starting from the future and working backwards. Specifically, closed-form expressions for the advancement boundary can be found by analyzing the points in the error-time plane from right-to-left and then from top-to-bottom.In the following steps, we start with the trivial case where all errors have been fixed (*E* = 0, Step 1) and calculate the advancement boundary for *E* errors provided we have carried out the analysis for *E* − 1 errors already (Steps 2-5):

1. The survival probability is equal to one when all *E*_0_ errors are repaired. It is obviously optimal to advance with the cell cycle when there are no errors (*E* = 0). Therefore, we draw an advancement boundary line at *E* = 0 (Fig. 3 A). Next, we iterate the following Steps 2-5:
2. For states with *E* unrepaired errors with an advancement boundary line at *E* − 1, we consider what happens when we increase *t*, keeping *E* fixed.At some time 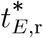, it may cease to be worth waiting for another repair event.This time is reached when the conditional repair probability times the gain in fitness for making one more repair, 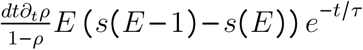, is outweighed by the loss of fitness if another repair does not occur (−*s*(*E*)*dt∂*_*t*_*e*^−*t*/*τ*^), depicted in Fig. 3 B. Equating these two terms and writing the survival probability in terms of independent contributions for each error, we obtain,

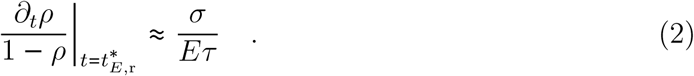

**Fig. 3:**
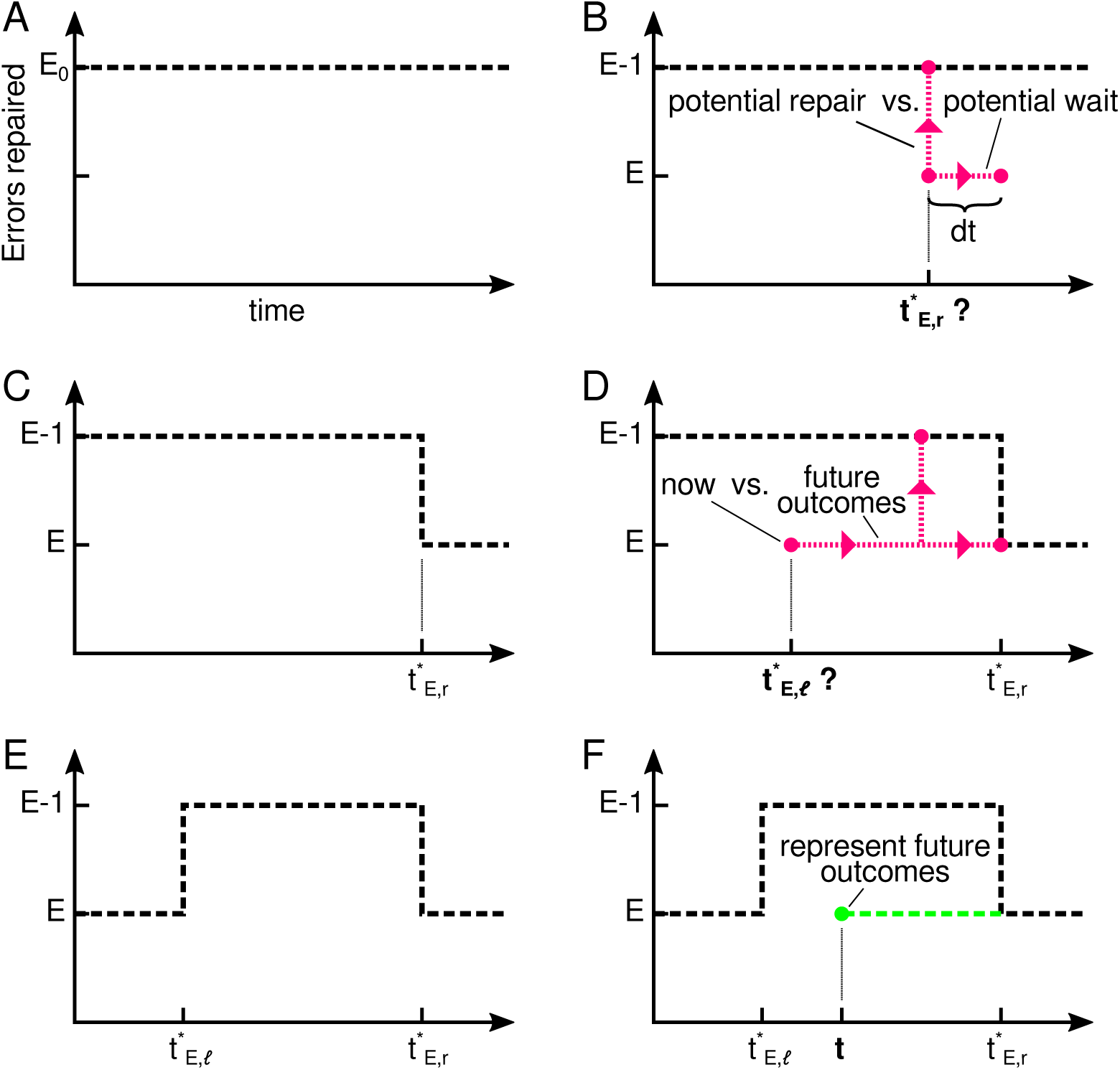
Steps for computing the optimal advancement boundary.

This relation allows the right advancement boundary 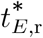 at *E* (Fig. 3 C) to be computed. (See Supplementary Information for additional details.)

Steps 3-5 of the computation and further details are presented in the Supplementary Information. In brief, in Step 3, after computing tright advancement boundary 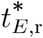 in Step 2, we find whether the wait up to the right boundary is preceded by any times 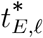 that should also trigger cell cycle advancement for the same number of *E* errors (Fig. 3 D, E). In Step 4, we find whether there are a series of such advancement-boundary–wait segments before (to the left of) 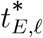, repeating Steps 2 and 3 for the same nu ber of errors *E* from right to left (Fig. S3 A). Before moving down in the diagram and repeating this analysis for *E* + 1 errors in the recursive computation, the time points inside the gapsait times) between 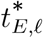 and 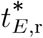 must be filled with predictions of the expected number of progeny if these time points are reached (green dashed line in Fig. 3 F). This is because the formulas in Steps 2-4 rely on the survival probability for cell cycle advancement with *E* (*t*) − 1 errors but if there is no advancement boundary at *E* (*t*) − 1, the expected number of progeny is the more general quantity to replace *s* (*E* − 1) in those formulas. So, in Step 5, the gaps between advancement boundaries for *E* errors are filled with the expected number of progeny once a particular time point in the gap is reached (Fig. S3 B).

The strategy encompasses two possible actions: i) wait for more potential repairs or ii) advance because waiting any longer brings down the expected offspring number. The reasons for advancement (ii) can be subtly different: At a border to the right of a waiting episode as in Fig. 2 E, any further waiting is disadvantageous (waiting is hopeless); arriving at a left border, before it is optimal to wait, a number of repairs must have been performed quickly, and any further improvements are unlikely to occur soon enough to be worth the wait (cash in a lucky draw); advancement at a top border can be interpreted either way.

The strategy is optimal by construction. Given that we start in the future and work backward, at each step we calculate the optimal decision based on having calculated the best decision for all accessible future time points.

### Experimental results

#### Experimental system

To apply the theory, we created genetically mutated budding yeast strains. To avoid artificially creating templates for DNA repair by homologous recombination, we minimized the duplication and introduction of extraneous DNA, including markers, by using the URA3-insertion/5-FOA-pop-out method throughout.

We deleted the Start cyclins CLN1,3 and replaced the CLN2 promoter by the MET3 promoter. This allows cells to be blocked in the pre-Start growth (G1) phase prior to DNA replication by adding methionine to the growth medium (+Met) or to be released to start a new cell cycle by removing methionine (-Met).^50^ Blocking cells in G1 prior to DNA replication, where haploid yeast cells only have one of each chromosome avoids efficient template-based repair mechanisms based on the duplicate chromosome and allows the controlled generation of DSBs.

We further integrated a GAL1pr-HO construct in the genome. The enzyme Ho creates a DSB with a 4-nucleotide 3’ overhang^40^ at locations in the genome where we inserted the short (30 bp) Ho recognition sequence. Ho has a natural cut site at the mating type locus (MAT), which we abolished in cells of mating type *α* by 10 synonymous mutations in the alpha1 gene (MAT*α*-syn). Ho is widely used to create DSBs to study checkpoints, override, and DNA repair.^40^

To detect DSBs and their repairs in single cells under the microscope or by flow cytometry, we devised a simple trick: We inserted an HO cut site (HOcs) and a destabilized version of the yellow fluorescent protein gene yEVenus between the constitutive yeast promoter ADH1pr and the non-essential ADH1 gene, creating an ADH1pr-HOcs-yEVenus-ADH1 fusion (Fig. 4 A) in place of the genomic ADH1pr-ADH1. (We refer to the yEVenus protein as YFP to simplify the figure labels and the text.) When the HO cut site suffers a DSB, yellow fluorescence should be low; when the DSB is repaired, yellow fluorescence should turn back on. Among other advantages, this system allowed us to induce Ho briefly in galactose medium, switch to the preferred carbon source glucose, and focus on cells with DSBs based on their fluorescence levels.

**Fig. 4:**
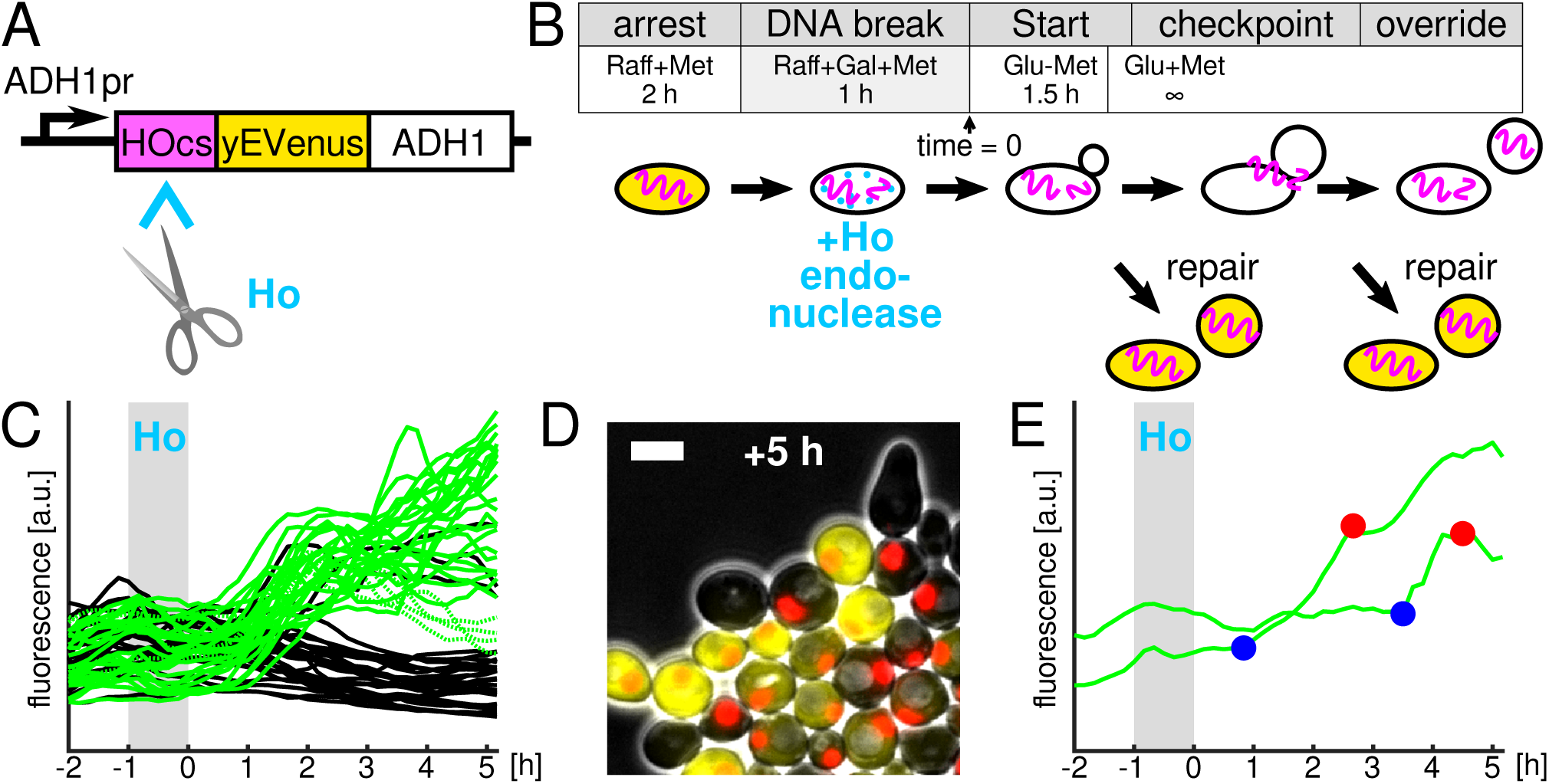
The ADH1pr-HOcs-yEVenus-ADH1 DSB sensor reports the presence of a DSB. A: Design of the sensor. B: Basic experimental protocol illustrated with clnΔ MET3pr-CLN2 GAL1pr-HO ADH1pr-HOcs-yEVenus-ADH1 strain. Raff = raffinose, Gal = galactose, Glu = glucose. C: YFP time courses in cells which additionally carry the cdc5-ad mutation. Green/black: cells that did/did not complete nuclear division. Four dead cells indicated by dotted lines. Switch to Glu-Met occurred at time 0 h. (n = 75) D: Cells 5 hrs after media switch to Glu-Met. Nuclear marker Htb2-mCherry in red. Scale bar: 5 *µ*m. E: Examples of YFP time courses for cells that divided nuclei late but seemed to repair the DSB early. Time point of rise in fluorescence indicated by a blue circle; nuclear division indicated by a red circle.

Thus, the basic genotype for all experiments described below is cln1-3Δ MET3pr-CLN2 GAL1pr-HO ADH1pr-HOcs-yEVenus-ADH1 HTB2-mCherry MAT*α*-syn, and only modifications of this strain are highlighted explicitly.

#### DSB repair distributions

A key input to the checkpoint theory is the repair probability distribution (*ρ*(*t*) in Eq. (2)). To quantify the timing of DSB repairs, we used the protocol depicted in Fig. 4 B: i) cells were arrested in G1 phase to ensure that only one copy of each chromosome was present, ii) a break was induced between ADH1pr and yEVenus by Ho, iii) the cell cycle was restarted, and iv) a new cell cycle was prevented. Step (iv) allowed us to watch cells for many hours under the microscope by preventing intact cells from dividing and overcrowding the field of view. This was necessary to detect late, rare repair events. Furthermore, in order to use nuclear division as an indicator of DSB repairs, we performed experiments with cells carrying the well-characterized cdc5-ad mutation^2^, which prevents checkpoint override, so that nuclear division only occurred in the absence of DSBs.

To characterize the system, we performed single-cell fluorescence microscopy and analyzed the images with the convolutional neural network YeaZ^51^. At the switch to Glu-Met, all cells showed low fluorescence due to the induced DSB or low ADH1 expression in raffinose and galactose. Subsequently, fluorescence shot up in all cells that divided their nuclei (Fig. 4 C). Thus, only cells where the DSB was repaired (or had never been induced in the first place, discussed below) divided. This underscores the effectiveness of override suppression by the cdc5-ad mutation since we saw no YFP-negative (YFP-) cells that performed nuclear division. Some cells did not divide their nuclei but showed a jump in fluorescence, indicating a failure to complete the cell cycle even in the absence of a DSB. We investigated the four cells that divided but showed low fluorescence after 5 hrs in Fig. 4 C (dashed lines) and found that they did not grow noticeably, suggesting that they were dead. Thus, there was a clear gap in fluorescence levels between healthy cells that divided and those that did not (Fig. 4 C, D). Furthermore, we compared the time when fluorescence increased and when nuclear division occurred and found that in all cells with dividing nuclei, fluorescence increased earlier than anaphase (examples of large differences between the two shown in Fig. 4 E). Thus, the DSB sensor provided direct information about whether cells had a DSB or not, which complements information provided by nuclear division. In the cells observed, anaphase was always preceded by rising YFP levels but YFP could increase without anaphase, i.e., repair could occur but the cell cycle fail, nevertheless.

As a first estimate of repair times, we recorded the time from budding to nuclear division by fluorescence microscopy with 10 min time resolution. We verified that the cdc5-ad mutation neither affected repair timing nor repair efficiency negatively by comparing the division times of cdc5-ad cells with wild-type CDC5 cells in which YFP turned on, i.e., did not override (61% vs. 51%, Fig. 5 A, B).

**Fig. 5:**
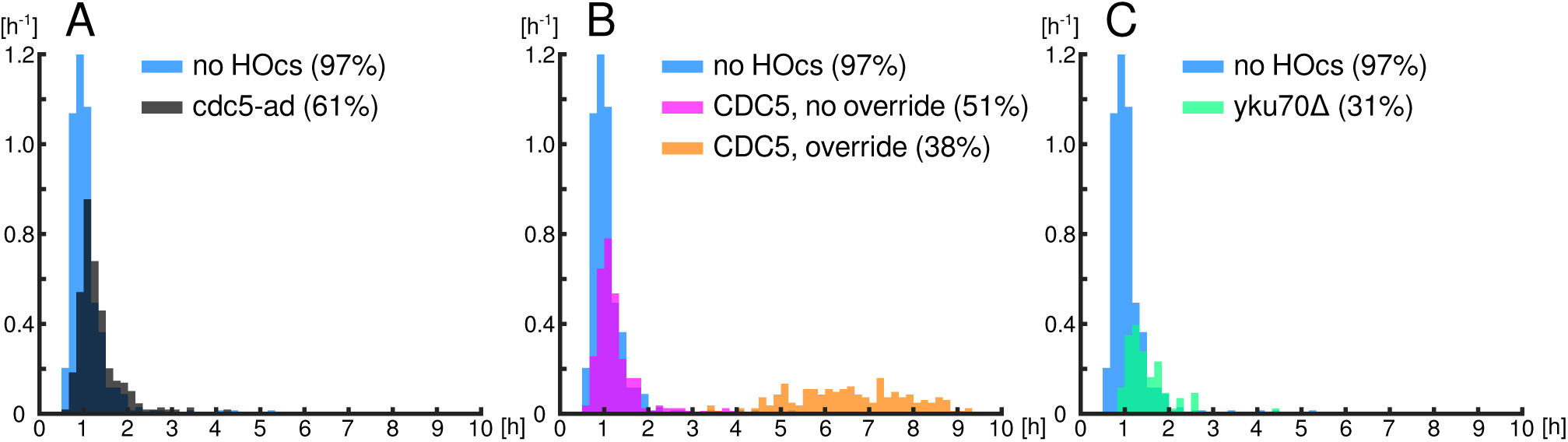
Rate of nuclear division with respect to budding scored by fluorescence microscopy. All divisions were accompanied by YFP increases except in panel B where dividing CDC5 cells could fail to turn on YFP (checkpoint override). n = 656 (cdc5-ad), 493 (wt-CDC5), 413 (no HOcs), 259 (yku70Δ).

A substantial fraction of cdc5-ad cells (39% = 100% – 61%) did not divide compared to 3% of cell cycles failing under these conditions in a no-cut-site control strain (Fig. 5 A). Thus, approximately 36% (= 39% – 3%) failed to repair the DSB. The fraction of cdc5-ad cells that divided (61%) (Fig. 5 A) did not simply represent inefficient cutting by Ho. This is because only 31% of cells that additionally carried the yku70Δ (=hdf1Δ) deletion, which blocks the non-homologous end-joining repair pathway^4^, divided, showing that 30% (= 61% – 31%) of cells repaired the DSB by a YKU70-dependent mechanism (Fig. 5 A, C). The fraction of dividing cells with the yku70Δ deletion is higher than in previous studies^38,45^, possibly due to differences in the DSB loci, e.g., level of transcriptional activity around the cut site. As expected, in the 10, one-tailed) higher than when there was no cdc5-ad population (Fig. 5 A) there were cells that divided late, i.e., later than 2 hours after budding, 6.1%, which was significantly (p = 2 · 10^−3^ cut site, 2.7%, presumably representing late repair events. However, most YKU70-dependent repairs (NHEJ) clearly took place very quickly (Fig. 5 A, C).

The long tails of the distributions observed by microscopy (Fig. 5) are potentially critical to understanding checkpoint override. However, statistics on the rare events that make up the tail were by their nature difficult to ascertain accurately by single-cell microscopy. Thus, we took advantage of the strong fluorescence signal of the ADH1pr-HOcs-yeVenus-ADH1 reporter to analyze cells by fluorescence-activated cell sorting (FACS) (Figs. 6 A, S4). At +4 hrs after the switch to Glu-Met (Fig. 4 B), > 10^6^ YFP-cells were isolated. To avoid contamination by YFP+ cells, we chose the gates for the FACS conservatively, taking into account the width of the distribution of fluorescence in a no-cut-site control strain (Fig. S4 C). (The yku70Δ results, discussed below, indicate that stray low-fluorescence but uncut cells were dead or negligible.) Then, from this population, multiple batches of 50 000 YFP- cells were sorted and plated on Glu-Met plates every 2 hours. On Glu-Met plates, cells could generate colonies if they repaired the DSB eventually since the absence of methionine reactivated the MET3pr-CLN2 construct and allowed new cell divisions. The double sorting, once at 4 hrs and once before plating, served to minimize the possibility that sorting errors let bright (YFP+) cells slip through as YFP- cells. After 3 days, we counted the number of all colonies on the plates and determined the subset of YFP+ colonies. With yku70Δ cells, substantially fewer colonies emerged, showing that nearly all colonies represented YKU70-dependent repairs by non-homologous end joining (Fig. 6 B, Fig. S4 B), and not technical artefacts. The number of YFP+ colonies decreased rapidly between the 6 h to 12 h time points (Fig. 6 B), showing that some YFP- cells had repaired the DSB between sorting events, had turned YFP back on, and had left the population of YFP- cells between time points. We estimate that for a typical YFP- cell, it took approximately 30 min to leave the YFP- population if it repaired the DSB (Methods). The total number of colonies, that is, YFP+ and YFP- combined, did not drop appreciably between the 6 h to 16 h time points (Fig. S4 D); this shows that the precipitous drop in YFP+ cells with time was not because cells were generally dying during the long arrests. Because the number of YFP- colonies did not change substantially within a very long time span (6 h-16 h), it is plausible that these were cells that repaired the DSB early but which destroyed the DSB sensor in the process and became permanently YFP- cells.

**Fig. 6:**
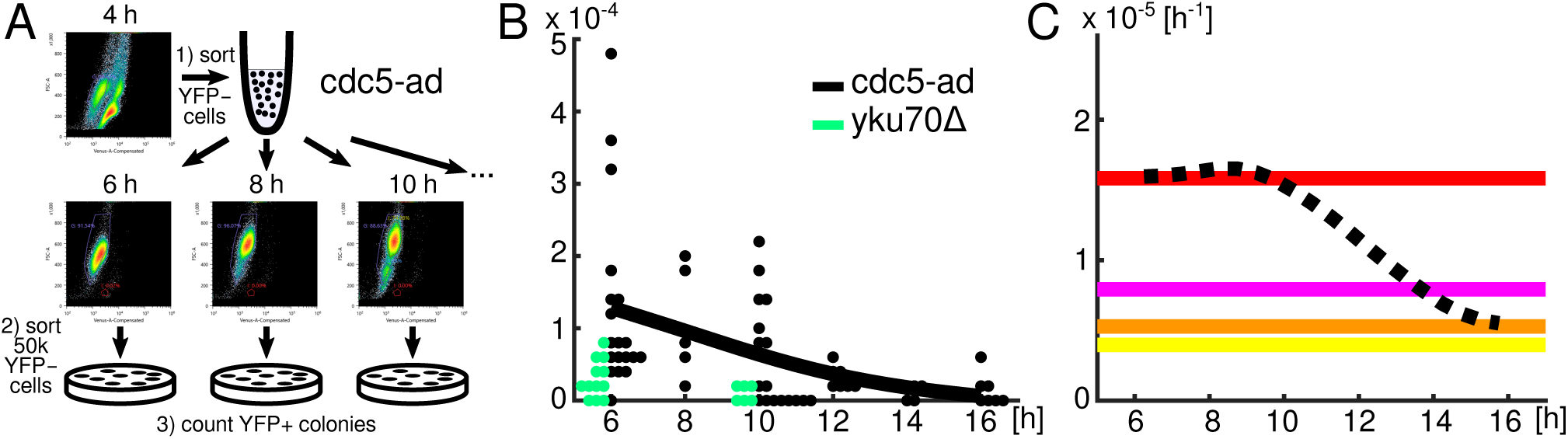
Measurements of late DSB repair statistics and survival rates after checkpoint override. A: Schematic of FACS experiments to measure the tail of the DSB repair time distribution. Horizontal and vertical axes on FACS plots are Venus-A and FSC-A, respectively. B: Measurements of the fraction of repaired cells (YFP+ colonies) compared to the number of cells plated (5 · 10^4^). Each circle represents the fraction of YFP+ colonies among 50k cells on one plate. Circles are stacked horizontally when replicas had the same numbers of colonies. The thick black line represents a spline fit to the mean values for cdc5-ad cells at each time point. C: The probability distribution function of repair (dashed line), which is the negative of the derivative of the fit in panel B. Horizontal lines represent *σ*/ (*Eτ*) for *E* = 1 (red), 2 (magenta), 3 (orange), 4 (yellow).

To extract the repair probability in an unbiased manner, we fit a spline through the mean fractions of repaired cells (Fig. 6 B). The negative of the derivative of the fit represents the conditional probability density function for DSB repair, *∂*_*t*_/*ρ* (1 − *ρ*), indicated by a dashed line in Fig. 6 C.

#### Statistics of survival after checkpoint override

Another crucial quantity determining the balance between risk and speed is the lethality of overriding the checkpoint with unrepaired damage (Eq. (2)). We measured the survival probability *s* (*E*) for one DSB (*s* (1) = *σ*) by repeating the experiment in Fig. 4 with wt-CDC5 cells. We plated CDC5 cells at the 6 h time point, just as cells were beginning to override the DNA damage checkpoint and after most pre-override repairs had taken place (Figs. 5 B, S4 E). Subtracting the survival probability of cdc5-ad cells, similarly sorted at +6 h to further remove the contribution of pre-override repairs, we arrived at the mean survival probability *σ*, which is represented, after dividing by the cell cycle time *τ* = 90 min log 2 by the red horizontal line in Fig. 6 C.

#### Quantitative experimental validation of theoretical predictions

We begin with a comparison of theory and experiment for one DSB (*E* = 1). The red (*σ*/ (*Eτ*)) and dashed lines (*∂*_*t*_*ρ*/ (1 − *ρ*)) overlap very well in the time window 6-9.5 h (90% or 95% bootstrap confidence intervals: *∂*_*t*_*ρ*/ (1 − *ρ*) is within 25% or 33% of *σ* /*τ*, respectively, in the 6-10 h window, see Methods). Remarkably, the measured override time in Fig. 5 B is 7.4 ± 1.4 h (mean ± STD) or 7.3 ± 1.3 h (median, 20th to 80th percentile) (n = 188). (Note that the microscopy results, e.g., in Fig. 5 B, show bud-to-nuclear-division times, to which the time from +0 until budding has to be added for each cell (≈50 min) to compare to FACS results.) Our measured override times for 1 DSB agree with the override time of ≈8 h reported in multiple past studies.^1,2,7^ Thus, the optimal checkpoint theory explains for the first time the mean as well as the broad width of the override distribution for 1 DSB.

We wished to make new, untested predictions. Assuming that each DSB reduces the survival probability independently, *s* (*E*) = *σ*^*E*^, we predicted the optimal override time for multiple DSBs using Eq. (2). The repair probability distribution *∂*_*t*_*ρ*/ (1 − *ρ*) intersects *σ*/ (*Eτ*) at approximately 13.8 h for *E* = 2 DSBs and 16.2 h for *E* = 3 DSBs. A linear extrapolation of the last half hour of *∂*_*t*_*ρ*/ (1 − *ρ*) intersects *σ*/ (*Eτ*) at 19.1 h for *E* = 4 DSBs.

To compare to our theoretical predictions and to revisit past experimental results, we inserted additional DSB cut sites at the URA3 locus or in the promoters of DLD2 or MIC60. URA3 was chosen for comparison with previous work^7^. DLD2 and MIC60 are in long (≈10 kB) regions of the yeast genome which can be deleted without affecting viability.^52^ Measurements were performed as before by single-cell timelapse microscopy where budding and anaphase were scored. To focus just on checkpoint override, we modified the protocol in Fig. 4 B and made it similar to previous studies^1,2,7^: We turned GAL1pr-HO on continuously in Gal-Met medium instead of Glu-Met after time 0 to ensure that cut sites would be recut and could not be repaired.^38^ Thus, nuclear divisions indicated checkpoint override because error-prone repair, after which Ho can no longer recut, is too rare (probability ≈ 10^−3^) ^53^ to have affected our measurements. We assessed how much galactose medium affects override times and found a relatively small change to 8.9 ± 0.3 h (mean ± SEM, n = 53, bud to nuclear division) with the single DSB in ADH1pr-HOcs-yEVenus-ADH1.

Strikingly, our results differ from previous conclusions based on bulk culture measurements but agree with the predictions of the optimal checkpoint theory (Fig. 7 E): For two cut sites at ADH1 and URA3 we observed that 80% of cells overrode with bud-to-nuclear-division time 13.4 ± 0.4 h (mean ± SEM, n = 75) (Fig. 7 A). For the two cut sites at DLD2 and MIC60, the mean override time was very similar, 13.6 ± 0.4 h (mean ± SEM, n = 88, bud-to-nuclear division) (Fig. S5 A). For three and four cut sites, the fraction of overriding cells decreased (34% and 42%) while the proportion of clearly dead cells, e.g., cell wall ruptured, increased. Nevertheless, for cells which performed nuclear divisions, we measured the override times to be 16.6 ± 0.7 h (mean ± SEM, n = 38, bud-to-nuclear division) for 3 cut sites and 20.2 ± 0.6 h (mean ± SEM, n = 85, bud-to-nuclear division) for 4 cut sites (Fig. 7 B, C). Note that these shifts in override timing as a function of DSB numbers cannot be explained by the greater lethality of increasing numbers of DSBs since the above override times are solely computed from the observed checkpoint overrides, not averaged with non-overriding or dead cells. Thus, the match between theory and experiment is remarkably close (Fig. 7 E), especially considering the disparate nature of the experiments and the substantial cell-to-cell variability.

**Fig. 7:**
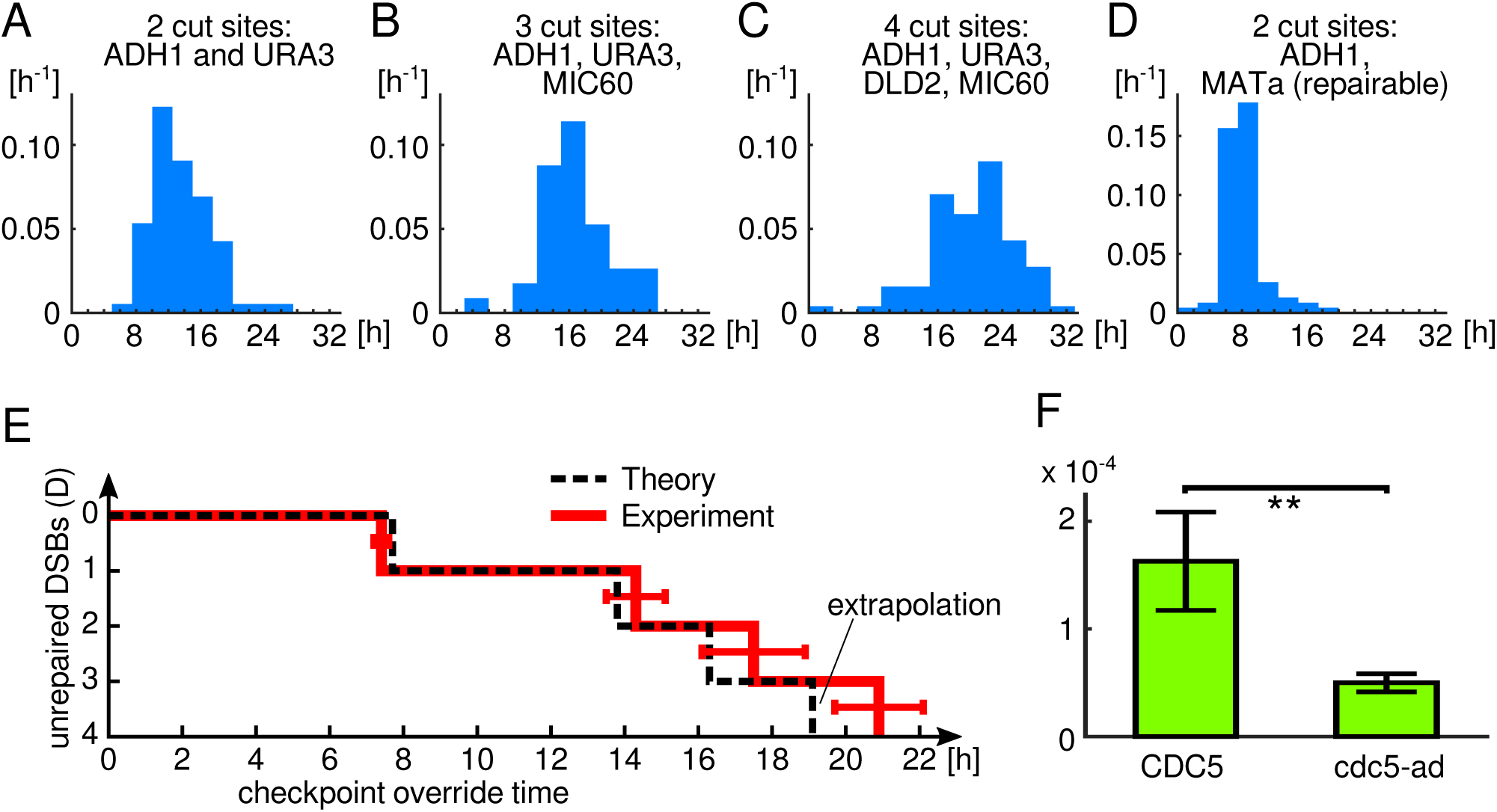
The optimal checkpoint theory predicts DNA damage checkpoint override times and checkpoint override is an advantageous strategy. A-D: Histograms of budding-to-nuclear-division probabilities for cells with multiple cut sites (n = 75, 38, 85, 92). E: Comparison of the optimal checkpoint theory with the experimental data. 50 min have been added to the means from the bud-to-nuclear-division histograms because the FACS results are with respect to time 0 in the experimental protocol. Horizontal red error bars indicate 95% confidence intervals. F: Direct comparison of survival probabilities of YFP- cells plated at the 6 h time point. Mean ± SEM shown. p = 0.0075, one-tailed. n = 7 (CDC5), 19 (cdc5-ad).

Thus far, we have compared our theoretical results for *E* = 1 DSB to experiments in which we ensured that cells had the DSB at the time of override by using the ADH1pr-HOcs-yEVenus-ADH1 sensor and focusing on YFP- cells. For *E* = 2, 3, or 4 cut sites, we ensured that DSBs were maintained throughout the experiment by continuous HO induction. (We repeated the *E* = 1 DSB experiment with continuous HO induction.)

Finally, we also wished to examine the predicted advancement boundary in a more complex way. We wondered, for example, whether the override decision could be confused by a decoy. To quantify more complex explorations of override decisions, we created a strain in which one DSB would be repaired efficiently and another DSB not. To accomplish that, we replaced MAT*α*-syn at the mating type locus, which could not be cut, by the wild-type MATa site in our DSB sensor (ADH1pr-HOcs-yEVenus-ADH1) strain. When MATa is cut by Ho, the DSB is repaired efficiently by homologous recombination based on the HML and HMR cassettes in the same chromosome.^54^ We returned to our original experimental protocol in which Ho is shut back off (Fig. 4 B). (We extended the induction window of GAL1pr-HO while cells were arrested in G1 to 3 hrs to ensure that in a large fraction of cells both MATa and ADH1-HOcs- YFP-ADH1 would be cut initially.) Control cells with only the MATa locus divided their nuclei very quickly after switching to Glu-Met (median: 75 min), showing that, indeed, MATa was repaired efficiently. However, in combination with the ADH1pr-HOcs-yEVenus-ADH1 sensor, YFP- cells overrode after 8.1 ± 0.2 h (mean ± SEM, n = 92, bud-to-nuclear division, Fig. 7 D), as expected for 1 DSB and unlike for 2 DSBs (Fig. 7 A). These results support our theoretical analysis, in which cells compute the optimal override decision continuously as they move in the error-time plane, and adjust their override time based on *E* (*t*), the present number of DSBs, as illustrated in Fig. 2 E.

#### Override is an advantageous strategy

The close match between predicted and measured override times (Fig. 7 E) suggests that checkpoint override may indeed be an advantageous strategy. However, this has not been shown directly in the past except with crippled DNA repair and maintenance genes^12,16,18^, leaving the question to what extent checkpoint override is relevant to wild-type cells open. In preliminary tests, we found a simple comparison of overriding (CDC5) to non-overriding (cdc5-ad) cells with one HO cut site under continuous HO induction inconclusive (data not shown). We speculated that most repairs occur well before checkpoint override (<6 hrs) and thus swamp the effects of checkpoint override (>6 hrs) on survival. Therefore, we compared CDC5 and cdc5-ad YFP- cells at the 6 h time point directly, using the protocol described in Fig. 6 A. We found a significantly higher probability of survival for CDC5 cells versus cdc5-ad cells (Fig. 7 F), showing that override is beneficial for cells with wild-type repair genes when they still have an unrepaired DSB late into the checkpoint arrest.

## Discussion

DNA damage checkpoint override follows predictable patterns. Here, we present the first theory of checkpoint strategies, which led us to discover these timing hierarchies.

Fundamentally, the theory is based on a population genetic analysis, which shows that the probability of producing live progeny multiplied by an exponential time penalty for arrest ought to have been maximized. In reality, this maximization would have been performed by evolutionary selection: Given two genetically encoded strategies, one closer to the optimum and one less optimal, the more optimal one would have produced more progeny and won out over evolutionary timescales – until a strategy even closer to the optimum would have displaced it in the population.

The theory is general in that it is compatible with both canonical models of population genetics, exponential growth as well as fixed-size populations. We demonstrated the latter by analyzing both the classical Wright-Fisher and Moran models. Furthermore, the theory is also general because it describes fundamental balances between risk, represented by repair and survival probabilities, and growth. It would presumably apply to other biological surveillance systems, including, for example, the spindle assembly checkpoint, even though in our presentation of the theory we anticipated the application to the DNA damage checkpoint. It can be extended straightforwardly, for example, to allow for sick cells.

We proved that according to the theory, a deterministic strategy is optimal. This may appear to be difficult to reconcile with the observed broad distribution of checkpoint override times (Fig. 5 B). However, the DSB repair probability is roughly flat for 6 h-10 h and then decays slowly (Fig. 6 C); thus, broad distributions of override times are compatible with a deterministic strategy since the optimal range is wide.

To solve for the optimal strategy, we mapped the problem of maximizing the fitness functional (Eq. (1)) onto the question of finding the optimal boundary for a random walk and derived closed-form recursive solutions. These can be calculated easily, e.g., Eq. (2). Simulations are not needed.

The theory establishes relationships between three independent quantities, i) the probability distribution of error correction, ii) the survival probability if errors are not fixed, and iii) the optimal timing of checkpoint override. The theory makes quantitative predictions for the override time as a function of the number of errors.

Experimentally, we built a budding yeast system which could be arrested in G1, a controlled number of DSBs could be induced, and the cells with DSBs could be separated from intact cells based on fluorescence levels from the DSB sensor. This system allowed us to switch to glucose as the carbon source after inducing HO in galactose. Switching HO expression back off in glucose permitted cells to repair the DSB. The fluorescent DSB sensor allowed us to eliminate repaired cells by flow cytometry, which pollute experiments with checkpoint-arrested cells because intact cells divide and overwhelm bulk measurements exponentially quickly. The alternative, keeping cells in galactose, was not an option because Ho would continuously recut the DNA (until the DSB would be repaired non-conservatively). We used continuous HO induction in galactose only to measure override times.

The experimental system allowed us to measure the parameters (i) and (ii) and the predictions (iii) precisely. The theory predicted the override times as a function of the number of DSBs strikingly well. The discrepancies between theory and experiment are less than 7% for ≤ 3 DSBs.

The predictions and experimental data are non-trivial. The override times vary by finite amounts as a function of the number of DSBs. These finite shifts were predicted well by the theory.

It is also interesting that the theory compelled us to reexamine the current view that override only occurs for 1 DSB and not 2 Ho-induced DSBs ^7^, which implies that checkpoint override is not a universal response to DNA damage. Instead, we found override to occur with up to 4 DSBs. We verified our override time measurements with two different DSB pairs, at ADH1 and URA3 or at MIC60 and DLD2 (Figs. 7 A, S5). We speculate that the wide distribution of override times for ≥2 DSBs (compare Figs. 7 A, S5 to Fig. 5 B), DNA re-replication after override, and slow post-override cell cycles hid the small fraction of post-override (1C DNA content) cells in bulk culture DNA content measurements at the time points 6 h, 12 h, 18 h, and 24 h in previous experiments^7^.

Finally, we measured the biological relevance of checkpoint override on cells without crippling DNA repair or maintenance. We filtered for cells that had not repaired their DSB shortly before override occurred and saw a significantly greater chance of survival for override-performing (CDC5) cells.

The remarkably close match of experiment to a theory that is based on elementary assumptions suggests further applications to other biological surveillance systems.

## Methods

### Strains

We performed experiments in budding yeast strains of background W303. Genetic manipulations and crosses were performed by standard methods. The cdc5-ad mutation was cloned into our strain background from a strain given to us by Achille Pellicioli. A GAL1pr-HO plasmid was given to us by Eric Alani.

The basic genotype for all strains was cln1Δ0 cln2Δ0::MET3pr-CLN2 (promoter replacement) cln3Δ0::GAL1pr-HO ADH1pr-HOcs-yEVenus-ADH1 HTB2-mCherry::HIS5 MAT*α*-syn. The modifications to this strain are indicated in Table 1. After constructing the basic strain, we performed three backcrosses with our wild-type W303 strains to reduce the likelihood of mutations.

**Table 1:** Strains used

|  |  |  |
| --- | --- | --- |
| AS18 | - | rad5-G535R |
| ET47 | - | RAD5 |
| AS20.1 | cdc5-ad | rad5-G535R |
| ET45 | cdc5-ad | RAD5 |
| AS30 | yku70Δ0 | rad5-G535R |
| ET46 | yku70Δ0 | RAD5 |
| ET44 | ADH1pr-yEVenus-ADH1 (no ADH1pr-HOcs-yEVenus-ADH1) | RAD5 |
| AS31.9 | ura3Δ0::HOcs::KanMX | rad5-G535R |
| AS32.1 | ura3Δ0::HOcs::KanMX | RAD5 |
| ET32 | MIC60pr::HOcs DLD2pr::HOcs (no ADH1pr-HOcs-yEVenus-ADH1) | rad5-G535R |
| RD58 | MIC60pr::HOcs DLD2pr::HOcs (no ADH1pr-HOcs-yEVenus-ADH1) | RAD5 |
| RD53 | ura3Δ0::HOcs::KanMX MIC60pr::HOcs | RAD5 |
| RD54 | ura3Δ0::HOcs::KanMX MIC60pr::HOcs DLD2pr::HOcs | RAD5 |
| AS14.2 | MATa (no ADH1pr-HOcs-yEVenus-ADH1) | RAD5 |
| AS35.2 | MATa | RAD5 |

HOcs stands for the 30 bp sequence: TTCAGCTTTCCGCAACAGTATAATTTTATA.

We performed experiments with cells carrying the W303-specific rad5-G535R mutation ^55^ as well as with cells with a corrected RAD5 gene wherever we indicate both versions of the strain in the list above. Since we observed no systematic or noticeable differences between results with rad5-G535R and RAD5, we pooled the measurements for higher statistical certainty. All other experiments were performed with cells with a corrected RAD5 gene only.

### Microscopy

Microscopy experiments were performed with commercial microfluidic chips. Timelapse recordings were carried out with a 60x objective and a Hamamatsu Orca-Flash4.0 camera. The interval between images was 10 min except for experiments with ≥2 cut site where the time between images was increased to 15 min to decrease potential phototoxicity during the long arrests. The override time was recorded as the first time point at which the Htb2-marked nuclei separated. In microscopy experiments where we released cells from Gal+Met into Gal-Met (Fig. 7 A-C), there were sometimes 1-2 cells that divided immediately and then arrested in the next cycle. We excluded these if this first division occurred within the first 100 min.

### Image processing

For segmenting the microscopy images, we used the YeaZ convolutional neural network and GUI^51^.

### FACS

Fluorescence-activated cell sorting was performed with a Sony SH800S instrument. The protocol and all the gates are illustrated in Figs. 6 and S4.

The instrument was set to room temperature for sorting events. A 100 *µ*m microfluidic sorting chip was used. The purity level was set to ‘ultra purity’. As excitation/emission filters we used those for ‘Venus’.

Before each sort, cells were centrifuges at 900 g for 1.5 min, medium was removed to concentrate cells, and the cell culture was sonicated for 8 sec. This was done to speed up the sorting. At the 4 h time point, 1.0 − 1.8 · 10^6^ YFP- cells were isolated per biological replica, which took about 20 − 40 min. After sorting, cells were centrifuged, the sheath medium was discarded, and ≈5 ml SCD+Met medium was added. Cells were always kept on a nutator at 30 C between sorting events. The second sorting events at 6 h, 8 h, 10 h, 12 h, 14 h, or 16 h of 50k YFP- cells took 3 − 8 min. After the second sorting, 100 *µ*l of SCD-Met medium was immediately added to the isolated cells and each batch of 50k YFP- cells was spread on a different Petri dish with SCD-Met agar medium.

To estimate the time it took cells to escape the YFP- gate after repairing the cut site at ADH1pr-HOcs-yEVenus-ADH1, we noted that at the 4 h sorting event, YFP- and YFP+ cells showed fluorescence levels of 1700 ± 700 [au] (median ± SD) and 7500 [au] (median), respectively. The speed of increase of fluorescence per hour therefore was about 1500 [au]/h, which is more than twice the standard deviation of fluorescence values in the YFP- gate. Hence, we estimate the escape time for repaired cells from the YFP- gate to be about 30 min.

### Fit to data

In Fig. 6 B, we fit a ‘SmoothingSpline’ through the means at each time point using the Matlab ‘fit’ function with parameter ‘SmoothingParam’ set to 0.1. A spline was used to avoid any biases regarding the functional form, e.g., type of decay, we might have expected. The smoothing parameter was chosen without fine-tuning, thus, it only has one significant digit. It was chosen among such one-significant digit values because it was the highest such number (generating the least amount of smoothing) that removed the bumps from the fit.

### Statistical tests

All p value tests were one-tailed. For the confidence interval analysis of the repair probability versus *σ* /*τ* in Fig. 6 C, we picked 10^3^ random alternative means for each time point (6 h, 8 h, 10 h, 12 h, 14 h, and 16 h) by bootstrapping. We fit the same spline as described above through 10^3^ combinations of these means and computed the derivative. We calculated how close the closest 90% and 95% of these conditional repair probability densities were to the *σ* / *τ* line between the 6 h and 10 h interval, which gave us the values quoted in the text (25% and 33%, respectively).

## Acknowledgments

SJR thanks Fred R. Cross for fruitful discussions and generous support as well as Eric Alani and Achille Pellicioli for reagents. We thank Dr. Enrico Tenaglia for technical advice and help with strain construction.

## Author contributions

AS performed all measurements including by FACS and microscopy and was helped in the latter by VG. AS, RD, and SJR made the constructs and strains. AS analyzed the data and was helped by RD and SJR. SJR wrote the manuscript. SJR devised the theory and calculations. All authors contributed to reviewing and editing the article.

## Supplementary Information

### Supplementary theoretical results

#### Outcome options: survival or death

Focusing only on the possibilities that cells either die or survive was supported by our observations in all DNA break experiments that the surviving cells grew into colonies of roughly the same size at about the same speed. Thus, we did not observe two different colony sizes, which would suggest healthy and sick survivors.

Similarly, other researchers have scored viability by counting the number of colonies that have been formed from single surviving cells without noting sick colonies. ^17,38^

Galgoczy and Toczyski^12^ observed colonies growing at different speeds after exposing diploid cells that could not repair DSBs by homologous recombination (rad52Δ) to X-rays. Diploid cells can survive the loss of a chromosome and become aneuploid. Thus, sickness may have been a more common outcome under those conditions than for haploid cells.

Furthermore, in reality, sick cells would be outcompeted exponentially quickly in time and thus would be effectively dead except in special circumstances, e.g., spatial isolation or immediate post-arrest mating and random assortment of the chromosomes generating some healthy offspring. These more complicated scenarios may be considered for further developments of the theory but we did not find those to be necessary because of the excellent match between theory and experiment.

In any case, the theory can be easily extended by introducing additional parameters describing the statistics of the occurrence and degree of sick cells.

#### Additional information on the arrest penalty

In a fixed-size population of *N* cells reproducing according to the Wright-Fisher model^47^, the survival probability of an arrested cell is (1 − (1 − 1/ *N*) ^*N*^) per generation. This factor represents the probability that the arrested cell is not lost from the population due to drift, which would occur if the *N* − 1 other cells in the population were selected to reproduce *N* times by chance. The probability of survival decreases from 3 /4 for *N* = 2 to 1 − *e*^−1^ = 0.63… as the population becomes larger (*N* → ∞). Since in this model, the checkpoint-arrested cell either dies or survives and then rejoins the regular population dynamics as before the arrest, the expected number of progeny is reduced by *P*_1_ [1 − (1 − 1/*N*)^*N*^]^*t*/*T*^ for a checkpoint arrest of *t*/*T* generations.

In a Moran process, the survival probability due to random drift is (1 − 1/ *N*) for a checkpoint-arrested cell, which is the probability that the cell is not killed with probability 1/ *N*. However, in a Moran process, it takes *N* generations on average for each cell to have its turn in reproduction. Thus, adjusted for the smaller time steps in this model, the survival probability in the presence of drift is (1 − 1/*N*)^*N*^ per *N* generations in this model, which increases from 1/4 for *N* = 2 to *e*^−1^ = 0.37… for *N* → ∞. The expected number of progeny after checkpoint arrest is thus *P*_1_ (1 − 1/*N*)^*Nt*/*T*^.

Thus, in all models, the expected number of progeny of a checkpoint-arrested cell is reduced by the product of a temporal penalty for arrest, which is exponential in time, and a probability ^−^of proliferation. In exponential growth, the fitness penalty due to checkpoint arrest (2^− *t*/*T*^) is exponential in time with base 1/2. In the Wright-Fisher model, the base of the exponential penalty depends on the size of the population; in each generation, the probability of survival decreases by a factor of 1 − (1 − 1 /*N*)^*N*^. In a Moran process, the equivalent factor is (1 − 1 /*N*)^*N*^. Interestingly, while these processes are very different, the loss of expected progeny per generation due to waiting is roughly similar numerically.

We represent the time penalty by a factor *e*^−*t*/*τ*^, which applies in all three cases and where *τ* absorbs the logarithm of the base, that is, *τ* = *T* /log 2 for exponential growth, *τ* = −*T* /log (1 − 1 − 1 /*N*) ^*N*^ for the Wright-Fisher model, and *τ* = −*T* /log (1 − 1 /*N*) ^*N*^ for the Moran model.

#### Fitness: arithmetic vs. geometric mean

To illustrate that with random checkpoint arrests, the average of the offspring numbers is an appropriate fitness function, we consider the following simple exponential growth model:

1. Starting with one cell, every cell produces two offspring with probability (1 − *q*) in every generation.
2. With probability *q*, a cell suffers damage, which is either lethal with probability (1 − *p*) or which is repaired with probability *p* and delays reproduction by *g* generations.

The expected number of offspring ⟨offspring⟩ for this model after *G* generations is approximately:

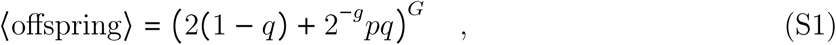

where 2 (1 − *q*) is the expected number of new cells due to normal divisions (model step 1) and 2^−*g*^*pq* is the expected number of cells which resolve the damage after arresting at the checkpoint for *g* generations (model step 2).

As the simulations of the model in Fig. S1 show, the above equation for the average number of offspring (⟨offspring⟩) captures the main peak of the distribution well. Thus, the average is a useful indicator for the behavior of the population. Given that the incidence of damage *q* is random and set by processes other than checkpoint control, we can only increase the average number of offspring by maximizing the product 2^−*g*^ *p*.

If, by contrast, we consider a fluctuating environment model in which a whole generation – not random cells – suffers damage with probability *q* which is lethal with probability (1 – *p)*, then in each generation, the chance of survival goes down by a factor of (1 – *q*(1 – *p*)). The survival probability thus goes to zero with time, and the average number of offspring after many generations would be dominated by unlikely scenarios where cells survive for long times and with many offspring. In these types of models the more appropriate fitness function to maximize would be the expected number of offspring for a typical (average) scenario, which leads to bet hedging.^49^

**Fig. S1:**
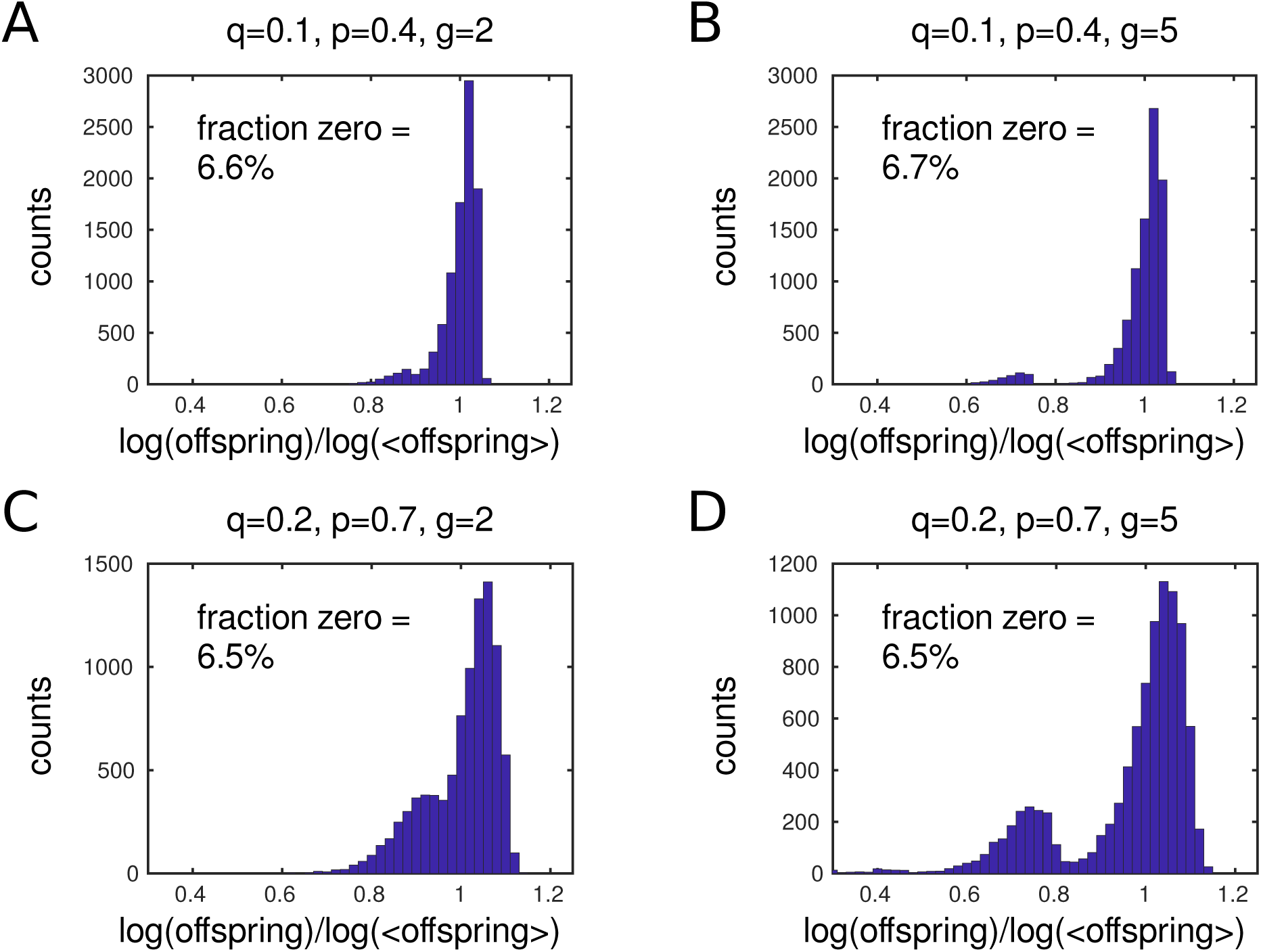
Distribution of offspring numbers for a simple population model of damage and checkpoint arrest, as explained in the Supplementary Information text, given the parameters listed on top of the histograms. As *g* increases, the first peak separates from the main peak. This small first peak represents trials where the very first cells in the simulation receive damage but arrest for *g* generations and repair the damage. The trials in which the first cell dies immediately (‘fraction zero’) are not indicated in the histogram because the logarithm of zero is undefined, but the fraction is noted in each panel. The simulation was run for twenty generations *G* = 20 and repeated 10^4^ times.

#### Additional information regarding the general fitness functional

To make the mathematical presentation simpler, we assume in Eq. (1) and in the subsequent calculations that if cells survive after cell cycle division, it is either always one of the two progeny that survives, in agreement with experimental data for DSBs ^56^, or always both progeny that survive. This assumption can be removed in the theory by introducing parameters for the relative likelihoods that one survives versus both survive.

#### Timer strategy

A simple checkpoint strategy could be to wait for a fixed time *t*′, and then advance with the cell cycle. Mathematically, this means setting *a*(*t*| {*t* _*i*_]) = *δ*(*t* − *t*′):

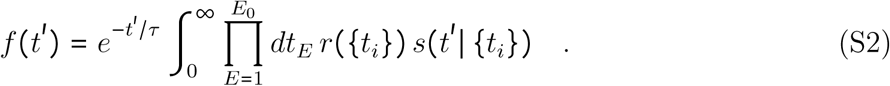

Assuming that each error is repaired independently according to the cumulative probability function *ρ*(*t*_*i*_) and each unrepaired error reduces the probability of survival by a factor *σ*, we obtain:

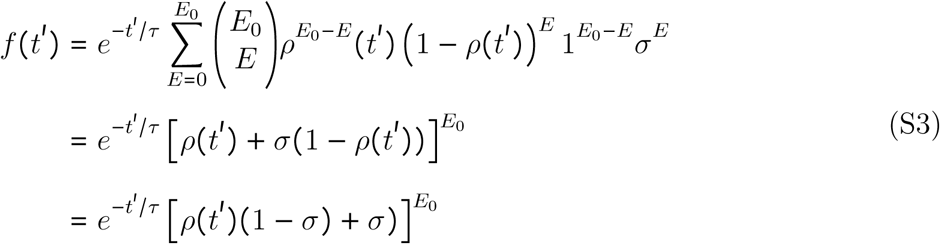

The best timer strategy is obtained by maximizing *f* (*t*′) with respect to *t*′,

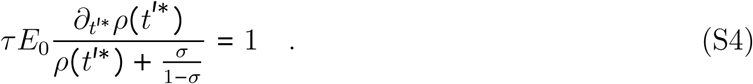

Assuming that repairs follow a Poisson process, 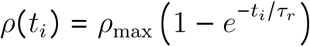, we obtain for the optimal waiting time in the timer model:

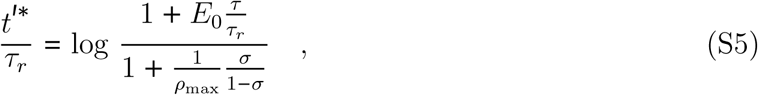

which implies that unless

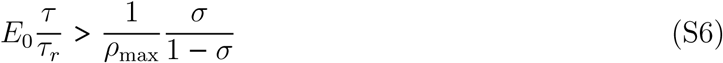

is satisfied, it is not worth waiting a fixed time for repair at a checkpoint at all (*t*^′*^ = 0), for example, if survival is too likely (*σ* close to 1).

#### Repair-threshold strategy

In an alternative model, a fixed amount (*E*_0_ – *E′)* of the total damage (*E*_0_) is repaired before continuing with the cell cycle. Here,

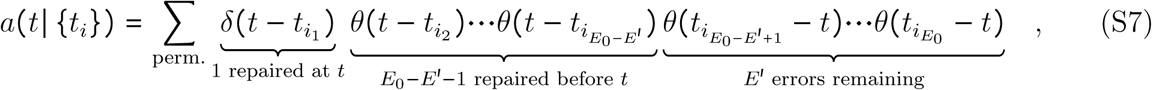

the cell cycle advancement probability, is equal to the sum of all permutations where damage *i*_1_ is repaired at time *t*, before which *E*_0_ − *E*′ − 1 errors were repaired, leaving *E*′ errors unrepaired.

Assuming as above that repair events are independent and survival for each error that remains at override is reduced by a factor *σ*, we obtain:

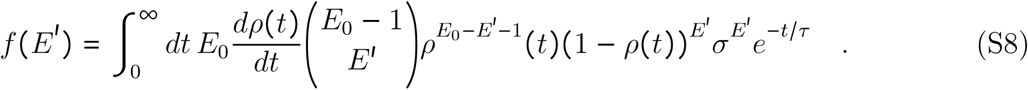

Assuming a Poisson process as above but with *ρ*_*M*_ = 1 to simplify the equations, we obtain:

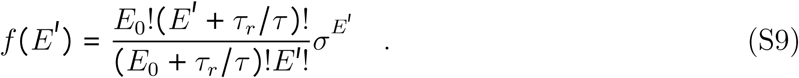

Maximizing *f* (*E*′) with respect to *E*′, we obtain:

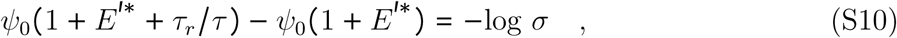

where *ψ*_0_ is the polygamma function of order zero.

The left-hand side is a decreasing function of *E*^′*^, so the larger the survival probability *σ*, the more errors *E*^′*^ can be tolerated for proceeding with the cell cycle.

*E*^′*^ is generally not an integer, and, therefore, *f (E*^′*^) has to be evaluated at the two nearest integers to *E*^′*^ to find the maximally tolerable amount of damage.

#### Optimal strategy is deterministic

The idea of the proof is as follows: To evaluate whether the optimal checkpoint strategy is stochastic or deterministic, we take a particular checkpoint strategy and increase or decrease the advancement probability *a (t*, {*t*_*i*_]) at one point (*t*, {*t*_*i*_]) in that strategy. Since the advancement probability must be normalized, that is, the integral or sum over all advancement probabilities must be equal to one, we also multiply all the other advancement probabilities that branch off from that point and that describe future advancement decisions by a compensatory factor. While satisfying the normalization constraint, we show that the fitness coefficient of the strategy *f* [*a*] can be increased unless the advancement probability density at that point was zero or infinity, that is, a Dirac delta (*δ*) function. A Dirac delta (*δ*) function advancement probability means that the checkpoint strategy is deterministic. Thus, only a deterministic strategy can be optimal.

To begin the proof, we assume that time is discretized with time points spaced Δ*t* apart to replace integrals by sums. Consider a checkpoint strategy with advancement probability *a (t*, {*t*_*i*_]) and a particular point in the strategy, that is, a damaged cell is arrested for time *t*, has *E* (*t*) errors, and has repaired *E*_0_ – *E* (*t*) errors at times {*t*_*j*_] which are smaller than *t*. The value of *a (t* | {*t*_*i*_]) of course only depends on *t* and the values of the repair times {*t*_*j*_] before *t*, since the repair times greater than *t* lie in the future. Therefore, *a*(*t* | {*t*_*i*_]) = *a* (*t* | {*t*_*j*_] holds. The contribution of this advancement probability to the fitness coefficient is:

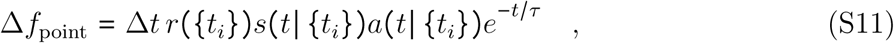

according to Eq. (1).

Next, we consider the later advancement probabilities 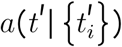 where *t*′ > *t* in which additional errors may have been repaired after *t* (the repaired errors before *t* are obviously the same *t*_*j*_), see Fig. S2 for an illustration. These advancement probabilities make the following contributions to the fitness coefficient:

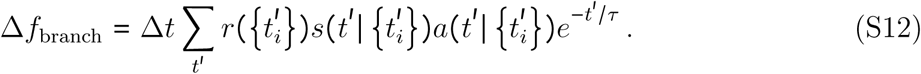

**Fig. S2:**
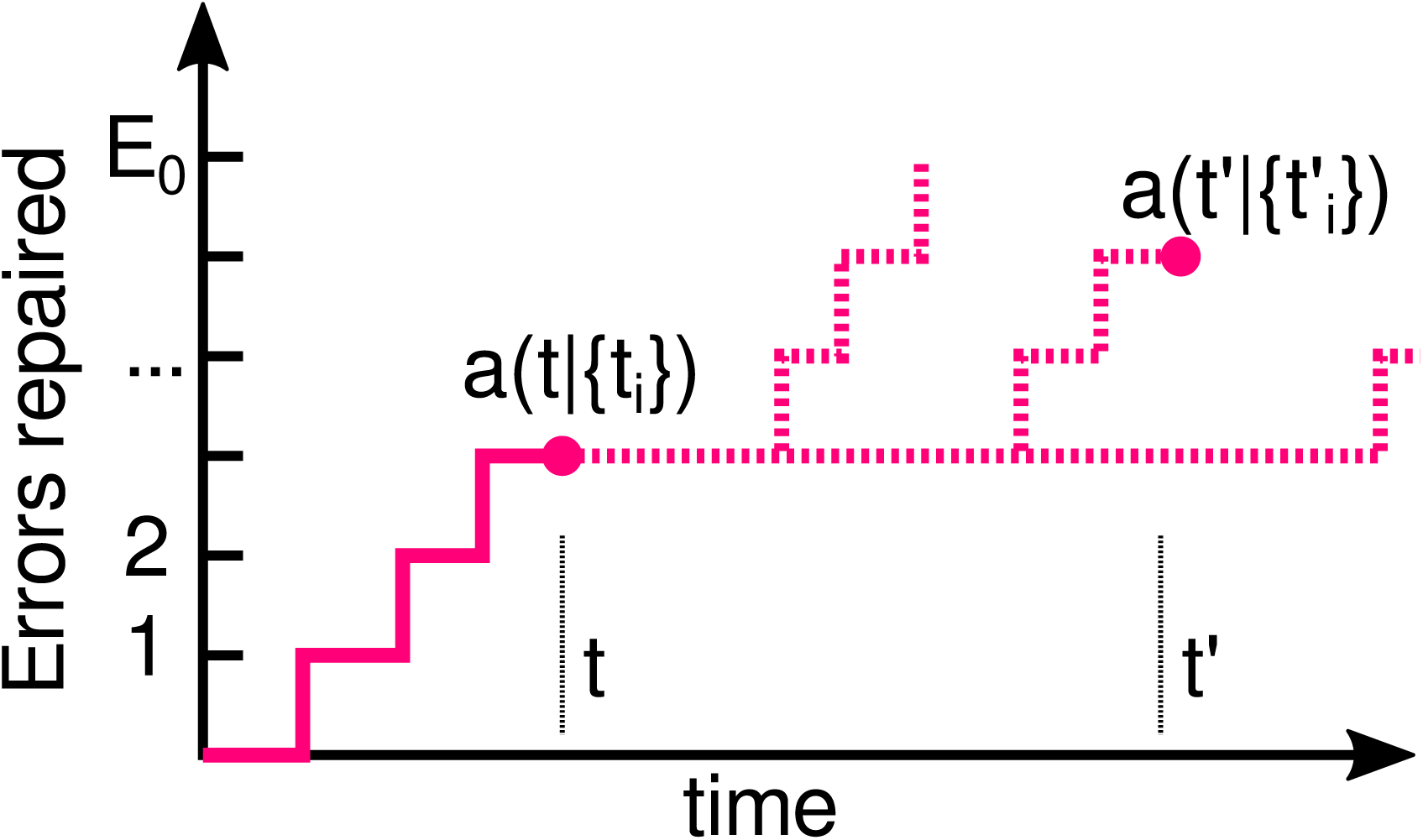
Illustration of trajectories in a checkpoint strategy. Solid lines indicate the realized trajectory (sequence of repairs) of the cell until time *t*. Dashed lines indicate possible future trajectories. The advancement probabilities *a*(*t*| {*t* _*i*_]) at time *t* and 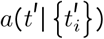 at a later time *t*^′^ are indicated.

Suppose that the advancement probabilities of the checkpoint strategy are normalized such that:

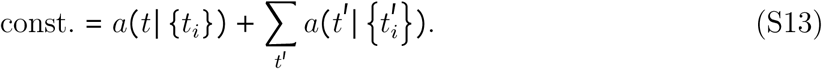

We create a new strategy in which we multiply *a*(*t*| {*t*_*i*_]) by a factor *α*, yielding *αa*(*t*| {*t*_*i*_]), and we also multiply each of the later advancement probabilities 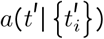 by *α*′:

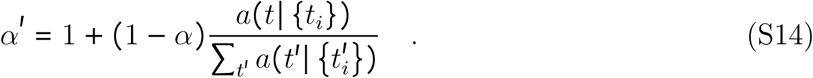

We see that in the new checkpoint strategy,

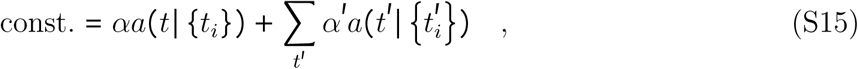

the advancement probability continues to be correctly normalized.

Plugging in the scaled advancement probabilities, we see that in this new strategy, the modified point contributes:

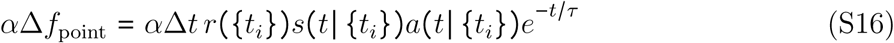

to the fitness coefficient and the contribution of the future points is:

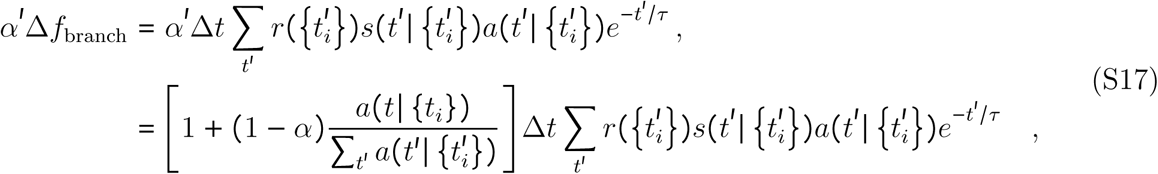

which is linear in *α*. Since the contributions to the fitness coefficient are linearly dependent on the parameter *α*,

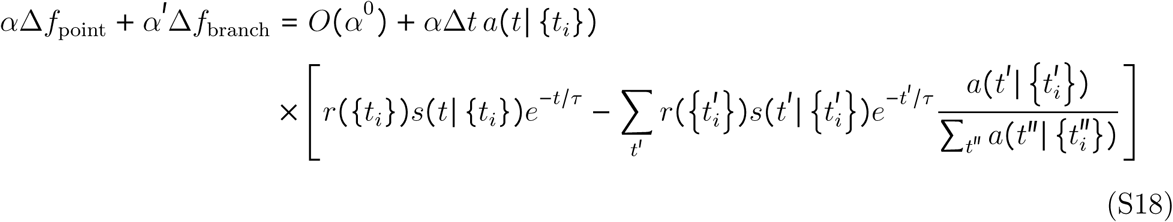

the fitness coefficient of this strategy could not be maximal, unless either *a*(*t*| {*t*_*i*_]) is zero or all future 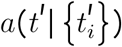 are zero. (Here, we assume that the slope of *α*, which is proportional to the term in square brackets in Eq. (S18), is not zero. This term is the difference between the contribution to the fitness coefficient if the advancement took place at *t* and the average contributions of all future divisions, which are weighted by their conditional advancement probability 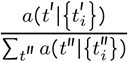. Generically, it will not be zero.)

This argument applies for every point in the error-time plane, such that at points where Δ*ta(t*, {*t*_*i*_]) is nonzero, it must be equal to one. In the limit Δ*t* → 0 as the discretization becomes smaller, *a (t*, {*t*_*i*_]) becomes a Dirac delta (*δ*) function.

#### Additional information regarding the optimal strategy

Here, we provide additional details for Step 2 (main text) for computing the optimal strategy and describe Steps 3-5 depicted in Fig. 3 D-F.

2. First, we provide more information regarding Step 2 in the main text. Equating the two terms discussed in Step 2, 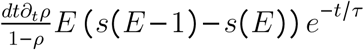 and −*s*(*E*)*dtδ_t_e^−t/τ^*, we obtain,

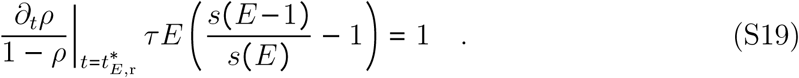

Next, we use *s* (*E*) = *σ*^*E*^ and only keep the linear term in *σ*, neglecting the *O(σ*^2^) terms. This is justified, for example, for an unrepaired DSB since the survival probability is very small (*σ* < < 1) (Fig. 7 D). This yields Eq. (2).

3. For the left advancement boundary at each *E*, we need to compare the instantaneous fitness *s* (*E*) *e*^−*t*/*τ*^ with the expected future fitness. To calculate the expected number of progeny due to waiting, one needs to take into account the possibility that another repair happens before reaching the right boundary at 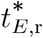 versus the possibility that no other such repair occurs (Fig. 3 D). This leads to the following condition:

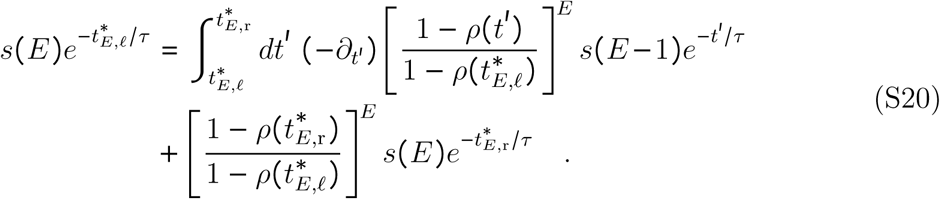

This sets the left advancement boundary 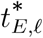 at *E* (Fig. 3 E). (The last two equations are related; Eq. (S20) simplifies to Eq. (S19) by taking the derivative with respect to 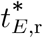 and setting 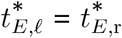.

4.It is, in principle, possible for there to be alternating segments of advancement boundary and waiting episodes (Fig. S3 A). Additional boundary points, if they exist, can be found by solving for the right-most 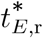, then the preceding 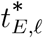, and then for a possible right boundary further to the left and so on.

5. Thus far, all calculations assumed that with one fewer error (*E* − 1), it is optimal to advance and that there is a continuous advancement boundary line, as for *E* = 0. This assumption explicitly entered the right-hand sides of Eqns. (2), (S19), and (S20) which refer to the survival probability if one more error is fixed (*s* (*E* – 1)) at the advancement boundary at *E* − 1. However, if at the previous step, which means at some time point *t* and for *E* − 1 errors, the optimal decision is to wait, it would not be correct to assess the consequence of reaching that point by the survival probability *s (E* – 1) since upon arriving at *t* and *E* − 1, the cell cycle should continue to arrest. So, to calculate the optimal checkpoint decision for *E* + 1 errors after the advancement boundary for *E* has been computed, for all times between the advancement boundaries, e.g., between 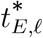 and 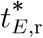 in Fig. 3 F, the expected number of offspring *f* _wait_ (*t, E*) needs to be computed. The formula for *f*_wait_ (*t, E*) has to be adapted to the specific situation but can be based on the following example:

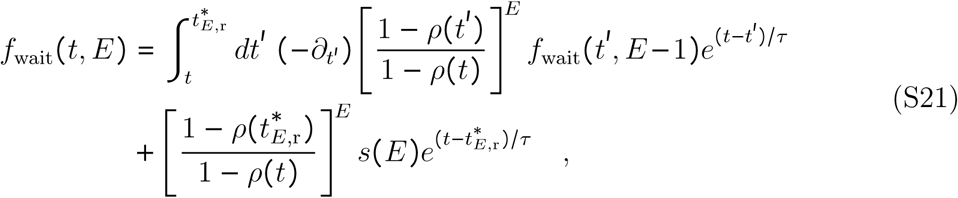

which describes a strategy in which the cell will wait at *E* − 1 (because there is also a *f*_wait_ (*t*′, *E* − 1) term on the right-hand side). *f*_wait_(*t, E*) replaces *s*(*E*) in the above formulas for calculating the advancement boundary for *E* + 1 errors (Fig. S3 B).

For a Poisson process 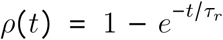, one can see by plugging into Eq. (2) that the boundaries are always horizontal, thus the optimal strategy reduces to the repair-threshold strategy. (The resulting condition, *τ* /*τ*_*r*_*D (*1 /*σ* −1) = 1, is not obviously equivalent to Eq. (S10), but we checked numerically, scanning a large range of parameters, that they both give the same optimal horizontal boundary.)

**Fig. S3:**
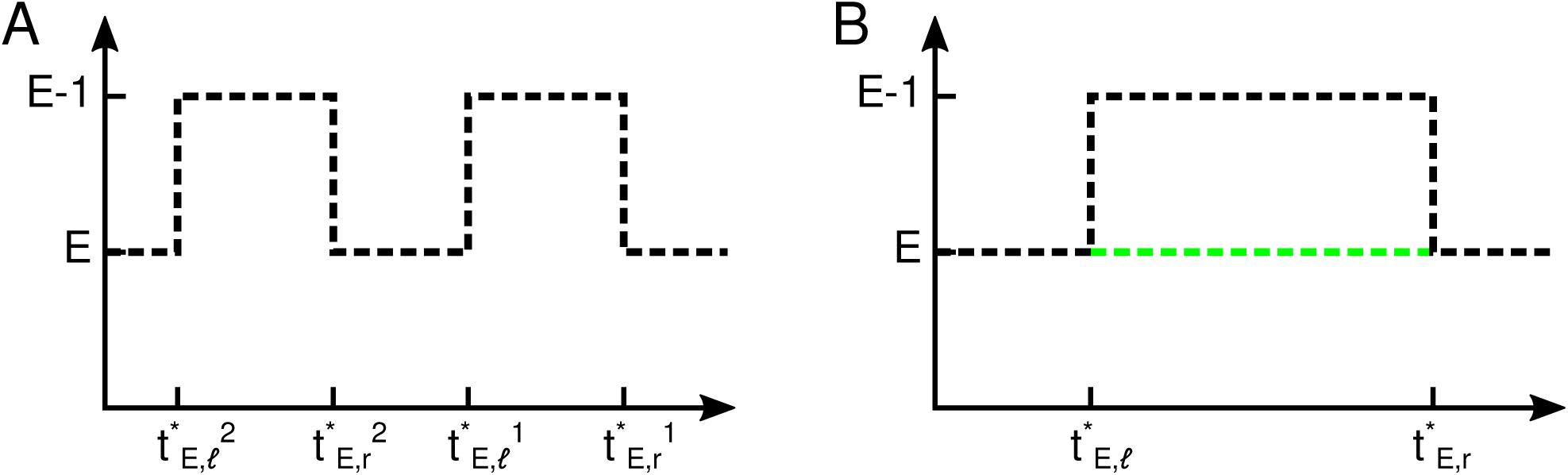
Additional plots to illustrate the recursive solution to the checkpoint optimization problem. A: Boundary with alternating advancement boundaries (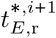 to 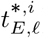) and wait segments (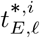 to 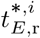). B: After all points for *E* have been analyzed, there are either advancement segments (black) or wait segments (green) with the expected numbers of progeny computed at each point of the wait segments. The calculations can then be repeated for *E* + 1 errors.

## Supplementary experimental results

### Fluorescence-activated cell sorting

**Fig. S4:**
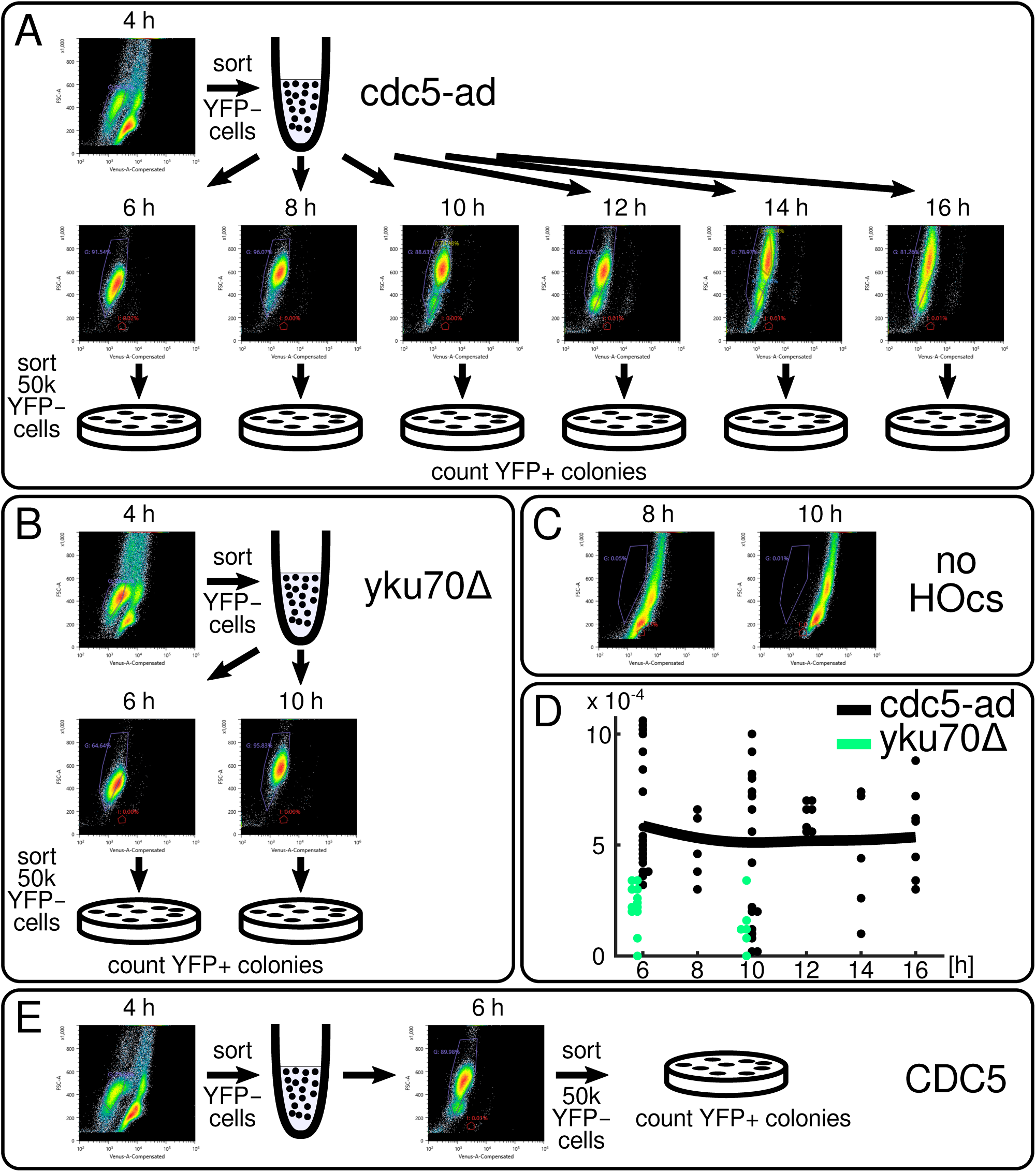
Supplementary data for the FACS experiments. A-C, E: FACS gates and protocols for the indicated strains. D: Total fraction of colonies (YFP- and YFP+ together) relative to 50 000 YFP-cells that were plated. Same experiments as shown in Fig. 6 B but only the fraction of YFP+ colonies is shown in Fig. 6 B.

### Multiple cut sites

**Fig. S5:**
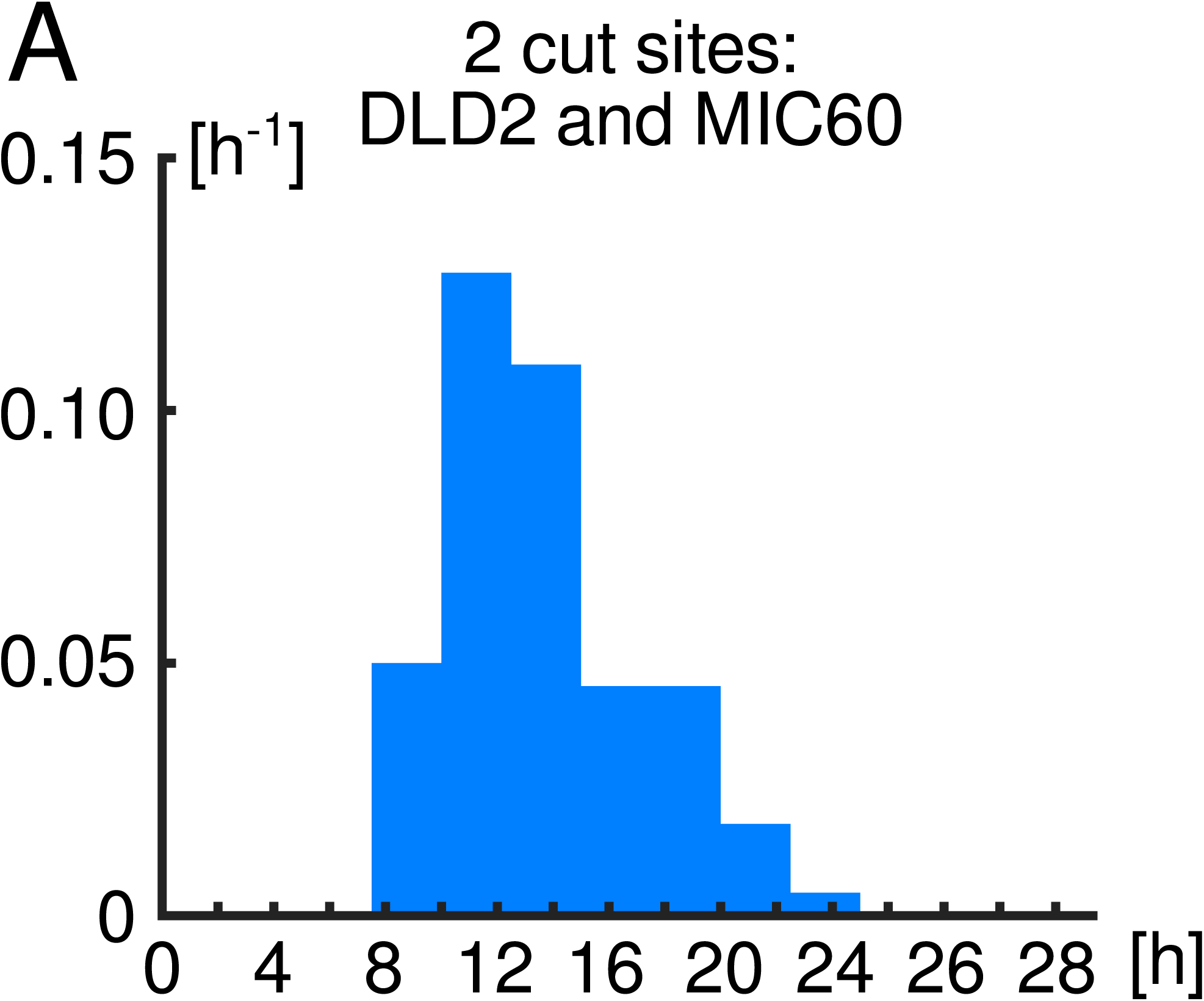
Histogram of budding-to-nuclear-division probabilities for cells with the two cut sites indicated. n = 88. Figure related to Fig. 7.

